# A blended genome and exome sequencing method captures genetic variation in an unbiased, high-quality, and cost-effective manner

**DOI:** 10.1101/2024.09.06.611689

**Authors:** Toni A Boltz, Benjamin B Chu, Calwing Liao, Julia M Sealock, Robert Ye, Lerato Majara, Jack M Fu, Susan Service, Lingyu Zhan, Sarah E Medland, Sinéad B Chapman, Simone Rubinacci, Matthew DeFelice, Jonna L Grimsby, Tamrat Abebe, Melkam Alemayehu, Fred K Ashaba, Elizabeth G Atkinson, Tim Bigdeli, Amanda B Bradway, Harrison Brand, Lori B Chibnik, Abebaw Fekadu, Michael Gatzen, Bizu Gelaye, Stella Gichuru, Marissa L Gildea, Toni C Hill, Hailiang Huang, Kalyn M Hubbard, Wilfred E. Injera, Roxanne James, Moses Joloba, Christopher Kachulis, Phillip R Kalmbach, Rogers Kamulegeya, Gabriel Kigen, Soyeon Kim, Nastassja Koen, Edith K. Kwobah, Joseph Kyebuzibwa, Seungmo Lee, Niall J Lennon, Penelope A Lind, Esteban A Lopera-Maya, Johnstone Makale, Serghei Mangul, Justin McMahon, Pierre Mowlem, Henry Musinguzi, Rehema M. Mwema, Noeline Nakasujja, Carter P Newman, Lethukuthula L Nkambule, Conor R O’Neil, Ana Maria Olivares, Catherine M. Olsen, Linnet Ongeri, Sophie J Parsa, Adele Pretorius, Raj Ramesar, Faye L Reagan, Chiara Sabatti, Jacquelyn A Schneider, Welelta Shiferaw, Anne Stevenson, Erik Stricker, Rocky E. Stroud, Jessie Tang, David Whiteman, Mary T Yohannes, Mingrui Yu, Kai Yuan, NeuroGAP-Psychosis, Dickens Akena, Lukoye Atwoli, Symon M. Kariuki, Karestan C. Koenen, Charles R. J. C. Newton, Dan J. Stein, Solomon Teferra, Zukiswa Zingela, Carlos N Pato, Michele T Pato, Carlos Lopez-Jaramillo, Nelson Freimer, Roel A Ophoff, Loes M Olde Loohuis, Michael E Talkowski, Benjamin M Neale, Daniel P Howrigan, Alicia R Martin

## Abstract

We deployed the Blended Genome Exome (BGE), a DNA library blending approach that generates low pass whole genome (1-4x mean depth) and deep whole exome (30-40x mean depth) data in a single sequencing run. This technology is cost-effective, empowers most genomic discoveries possible with deep whole genome sequencing, and provides an unbiased method to capture the diversity of common SNP variation across the globe. To evaluate this new technology at scale, we applied BGE to sequence >53,000 samples from the Populations Underrepresented in Mental Illness Associations Studies (PUMAS) Project, which included participants across African, African American, and Latin American populations. We evaluated the accuracy of BGE imputed genotypes against raw genotype calls from the Illumina Global Screening Array. All PUMAS cohorts had R^2^ concordance ≥95% among SNPs with MAF≥1%, and never fell below ≥90% R^2^ for SNPs with MAF<1%. Furthermore, concordance rates among local ancestries within two recently admixed cohorts were consistent among SNPs with MAF≥1%, with only minor deviations in SNPs with MAF<1%. We also benchmarked the discovery capacity of BGE to access protein-coding copy number variants (CNVs) against deep whole genome data, finding that deletions and duplications spanning at least 3 exons had a positive predicted value of ∼90%. Our results demonstrate BGE scalability and efficacy in capturing SNPs, indels, and CNVs in the human genome at 28% of the cost of deep whole-genome sequencing. BGE is poised to enhance access to genomic testing and empower genomic discoveries, particularly in underrepresented populations.

## Introduction

Genome-wide association studies (GWAS) have grown exponentially over the last 15 years, rapidly increasing in statistical power to enable the identification of hundreds of thousands of associations between genetic variants and human traits.^1^ While these discoveries have been facilitated in part by precipitous drops in sequencing costs, microarrays have been the primary technology used for GWAS to date because of their lower costs. However, by design, they have biased ascertainment of genetic variants; sites that are included on many GWAS arrays, such as the widely used Illumina Global Screening Array or Global Diversity Array, are most common in European ancestry populations. Previous work has shown that low-coverage sequencing is a cost-effective alternative that can more accurately capture genetic variants across the allele frequency spectrum for variants present in imputation reference panels.^2,3^ Low-coverage sequencing is especially useful in populations underrepresented in genomics, even compared to GWAS arrays that have been designed to reflect variation within those populations, such as the H3Africa GWAS Array.^3^ Both genetic data generation strategies are useful for evaluating the architectures of complex traits and associating common variants with them.

High-coverage genome sequencing, while more expensive, captures a more complete spectrum of genetic variation. Balancing its higher cost with the need for large sample sizes to achieve robust disease and trait associations, researchers often sequence only coding regions using an exome capture, typically to high coverage (∼60X), and supplement with GWAS arrays. This strategy is useful for prioritizing genes, as, to date, rare coding variants comprise most of the known variants that have interpretable functions and are therapeutically actionable.^4–6^ Combining exome sequencing with GWAS arrays enables researchers to glean analytical insights from both common variant GWAS and rare variant tests, including those that assess gene burden and those that evaluate associations to individual variants. Another recent study has shown the utility of conducting high coverage exome sequencing in parallel with low coverage whole genome sequencing to gain additional, non-imputable disease associated variants available to be evaluated.^7^ To increase the cost efficiency and scalability of a combined strategy, we recently developed a new blended genome exome sequencing (BGE) approach.^8^ BGE sequences the whole genome to at least 1-4X depth and the exome at ≥30X depth, reducing costs. Another benefit of BGE compared to disjoint exome plus GWAS array or low coverage sequencing on the same samples is the unified protocol which streamlines comparisons between imputed versus high-quality and -coverage coding variants across the allele frequency spectrum and thus enables improved QC. However, it requires the development of new computational approaches and pipelines as well as haplotype reference panels that can support genotype calling and refinement.

Here, we demonstrate the utility of BGE across diverse populations that have been underserved with traditional GWAS arrays by applying BGE at scale by sequencing samples from the Populations Underrepresented in Mental Illness Associations Studies (PUMAS) Project. PUMAS is an umbrella project consisting of multiple sub-studies, including: the Genomic Psychiatry Cohort (GPC) that primarily includes admixed African American and Hispanic/Latino populations from the US; the Neuropsychiatric Genetics in African Populations Psychosis (NeuroGAP-Psychosis) Study consisting of participants from Ethiopia, Kenya, South Africa, and Uganda; and the Paisa Study, consisting of participants from a recently admixed population in Colombia. We report high genetic data quality at a relatively low cost when applying the BGE method to sequence 53,448 samples from the above populations.^8^ To facilitate benchmarks against gold-standard data, we also applied BGE sequencing to 400 individuals enrolled with a family study design from the Simons Simplex Collection (SSC) provided from the Simons Foundation Autism Research Initiative (SFARI),^9^ that had previously been sequenced to high coverage to evaluate recall and positive predictive value (PPV) for exonic copy number variants (CNVs) and structural variants (SVs). We also evaluate imputation concordance of BGE using an orthogonal data generation method, standard GWAS arrays, in a subset of the same PUMAS participants; we stratified by local ancestry where applicable. To support broader adoption, we provide a resource of open-source scripts and analytical pipelines for conducting these analyses with BGE (see Software Availability), empowering other sequencing centers and analysts to apply this new strategy to cost-effectively improve variant discovery, especially in underrepresented populations.

## Results

### The BGE protocol balances cost with genotype quality across coding and non-coding regions

As described previously,^8^ we performed six iterative rounds of experimentation prior to implementing BGE at a large scale to determine the best blending ratio of Whole Exome Sequencing (WES) to Whole Genome Sequencing (WGS) and amount of sequencing required per sample to achieve the following objective: cost-efficiently maintaining >99% imputation R^2^ non-reference concordance for imputed variants with MAF >5% compared to existing 30x WGS. The first round tested 96 samples per lane of NovaSeqS4 (Illumina, San Diego, CA, USA) and nanomolar blending of 67% WES: 33% WGS. This ratio generated 29x WES and 1.5x WGS coverage on average per sample and resulted in >99% R^2^ concordance of BGE imputed variants in the exome, but <99% for whole genome variants in samples with lower coverage. In the subsequent experiments (rounds 2-6), we titrated the WES:WGS blending ratio and sequencing coverage to determine the optimal protocol for achieving >99% R^2^ concordance for calling both exome and genome variants.^8^ **Table 1** shows the blending ratios, amount sequenced, and resulting WES and WGS coverage across rounds.

**Table 1.** Iterative rounds of blended genome exome (BGE) development. WES = whole exome sequencing, WGS = whole genome sequencing. Per-sample genotype concordance between deep whole genome variants and filtered Haplotype Reference Consortium (HRC) imputed variants from the low-pass genome, as described previously.^8^

| Round | 1 | 2 | 3 | 4 | 5 | 6 - Scaling |
| --- | --- | --- | --- | --- | --- | --- |
| Blending ratio | 67% WES + 33% WGS | 67% WES + 33% WGS | 60% WES + 40% WGS | 40% WES + 60% WGS | 33% WES + 67% WGS | 33% WES + 67% WGS |
| Amount of Sequencing | 96 samples on 1 lane NovaSeq S4 | 48 samples on 1 lane NovaSeq S4 | 48 samples on 1 lane of NovaSeq S4 | 48 samples on 1 lane of NovaSeq S4 | 64 samples on 1 lane of NovaSeq S4 | 768 samples on 2 lanes of NovaSeq S4 |
| Coverage (mean) | 29x WES<br>1.5x WGS | 78x WES<br>2.55x WGS | 78x WES<br>2.5x WGS | 64.8x WES<br>3.7x WGS | 37.7x WES<br>2.37x WGS | 37x WES<br>2.5x WGS |
| % Exome bases $\geq 10x$ | 96.33 | 98.34 | 98.27 | 98.18 | 96.44 | 96.37 |
| % Exome bases $\geq 20x$ | 75.09 | 97.84 | 97.79 | 97.19 | 87.04 | 84.08 |
| Genotype concordance $R^2$ median (sd) | 0.988 (1.56e-3) | 0.991 (1.28e-3) | 0.990 (2.90e-3) | 0.995 (3.55e-2) | 0.993 (1.15e-3) | 0.993 (1.05e-3) |

To achieve optimal blending and sequencing depth, we found that 33% WES and 67% WGS for 64 samples/lane provided adequate coverage of each (30-40x WES and 1-4x WGS per sample) for calling variants, and >99% r^2^ concordance to 30x WGS common variants (MAF > 5%) for both the exome and imputed genome. All BGE blending ratios tested provided better R^2^ concordance to 30X WGS data than the Global Screen Array (GSA).^8^ We recommend using the following target metrics for BGE sequencing: 1) >9.5 Gb total bases/sample passing Illumina’s PF filter, 2) >90% of exome bases at 10X coverage, and 3) between 10-65% of reads on/near exome capture regions (within 250-500 bp of the probe). These deliverables provide adequate exome coverage for rare variant detection and genome coverage for common variant imputation.

### BGE quality control at scale on ancestrally diverse participants

With our recommended BGE protocol in place, we sequenced 53,446 individuals (**Table 2**) from an ancestrally diverse set of participants collected in the PUMAS Project (**Figure 1A-B**, **Supplementary Figure 1, Methods)**. Exome coverage differed based on DNA collection method, with blood samples from GPC and Paisa slightly outperforming saliva samples from the NeuroGAP (percent failure rates: GPC = 0.04%, Paisa = 0.13%, NeuroGAP = 0.48%, **Figure 1C**). However, across all cohorts, the percentage of samples failing to meet the exome coverage threshold was consistently less than 1%, with only 6 samples removed for having less than 1x average WGS coverage (**Figure 1D**). Overall, the exome portion of the BGE demonstrated robust performance, with mean call rates > 0.99, mean read depth > 30, and mean genotype quality > 39 across all cohorts (**Figure 1D**). We defined high-quality samples using the exome portion of the BGE and continental ancestry strata-based filtering, which retained 88% of the samples (**Table 2, Methods & Supplement**).

**Figure 1.**
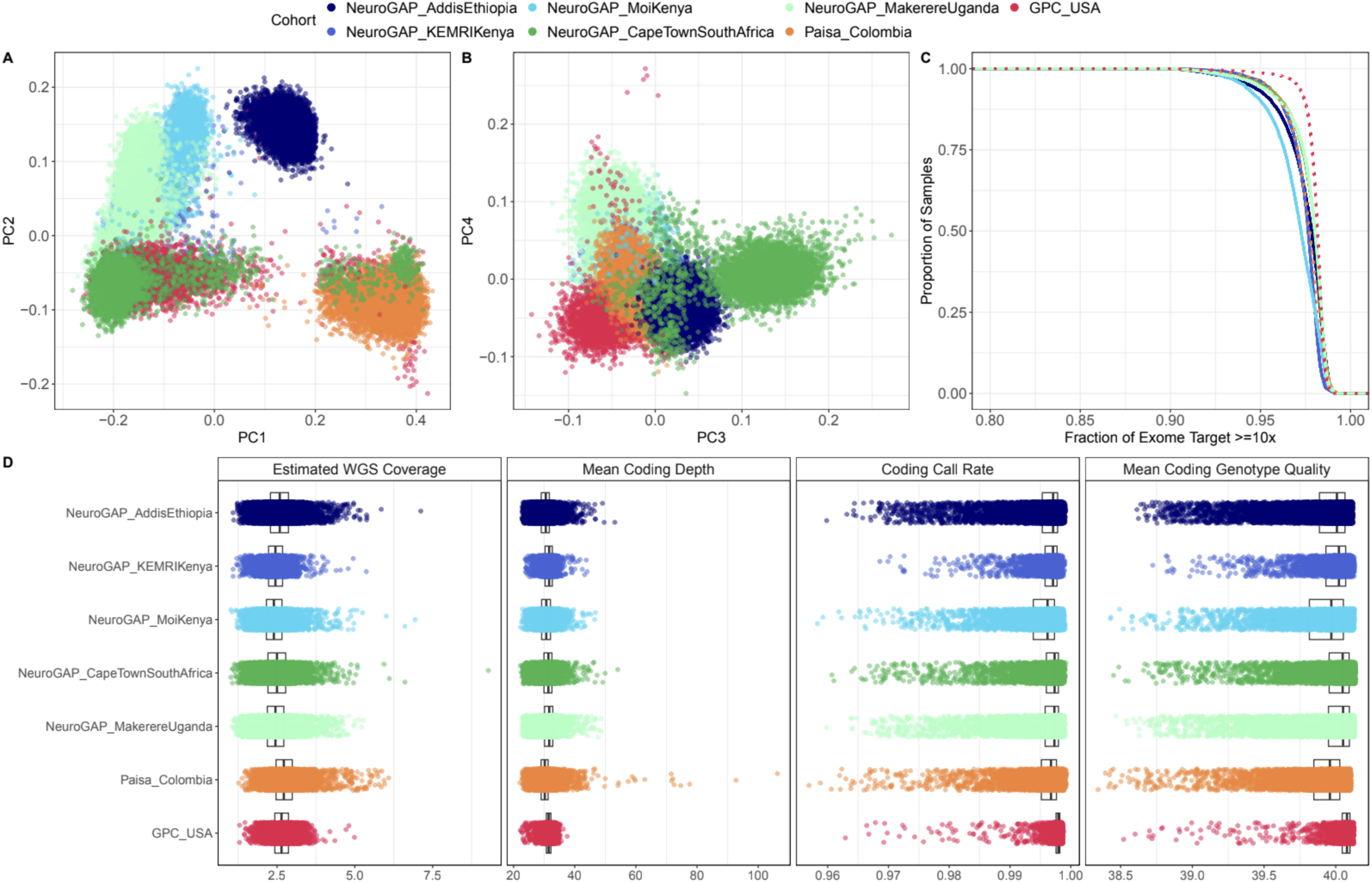
Expected ancestral diversity, coverage, and quality from BGE data at scale. A) Principal components (PC) 1 vs PC2 and B) PC3 vs PC4, C) Fraction of exome target covered with at least 10x depth stratified by cohort and collection method. Solid lines indicate saliva collection (NeuroGAP) and dashed lines indicate blood collection (GPC and Paisa). D) Estimated mean WGS coverage, mean coding depth, mean coding call rate and mean coding genotype quality.

**Table 2.**
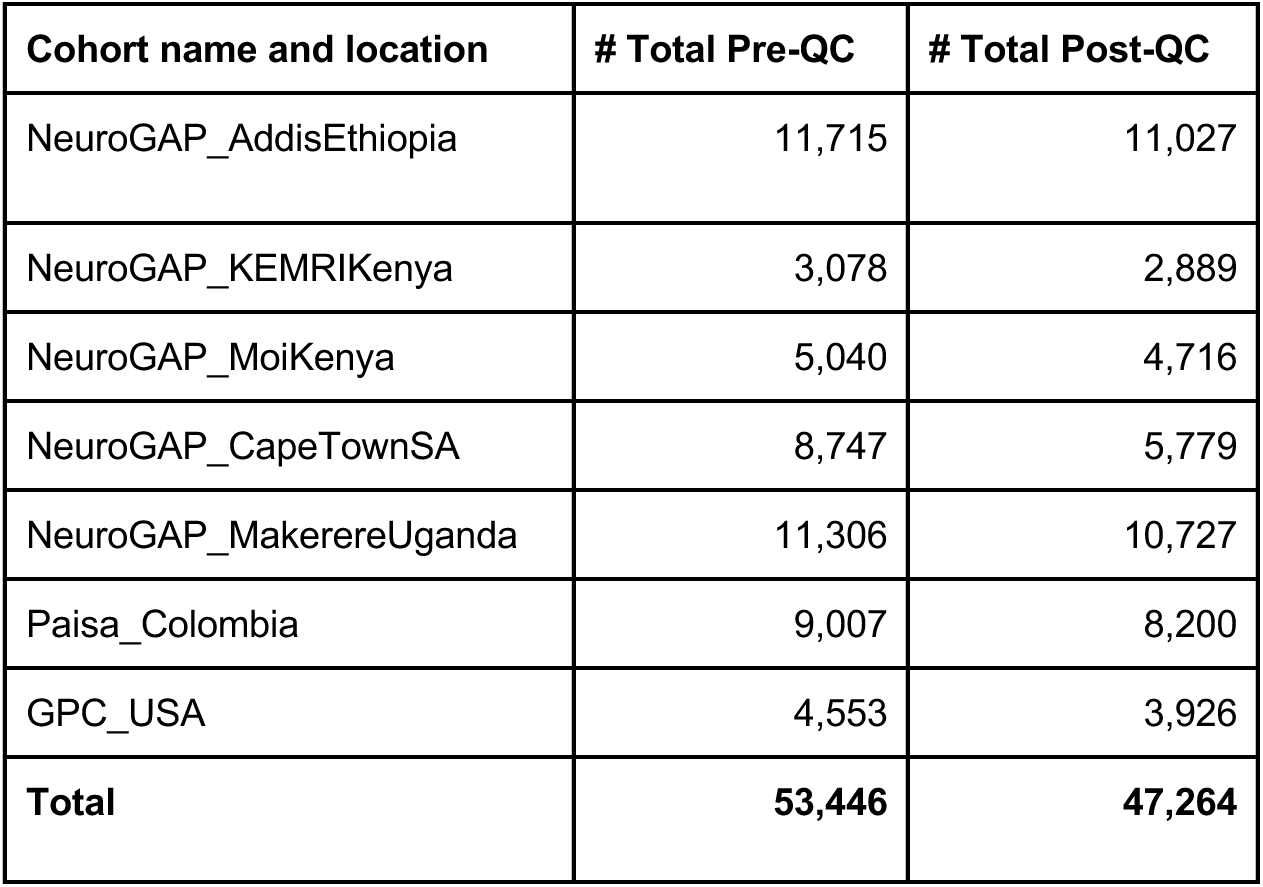
Cohorts included throughout analyses. Quality control includes filtering on WES and WGS coverage, genetic ancestry, outlier filtering on sample quality metrics, and checking for discrepancies between genetic sex and reported gender.

### Accurate copy number variant (CNV) and structural variant (SV) discovery in BGE data

We examined the potential of BGE for detecting CNVs, focusing initially on the higher coverage exome sequences. CNV discovery from WES data has traditionally been challenging due to the highly variable exome capture efficiency between different capture kits and sequencing centers, as well as the complexities introduced by using read-depth information to infer copy state from short-read sequencing data. These factors have most often resulted in restricting exome analyses of SNVs and indels, despite the considerable value of capturing and predicting the functional impact of CNVs that can alter gene dosage and/or disrupt normal gene function. Recently, members of our group have published the GATK-gCNV algorithm^10^, which is a read-depth based method built on a hierarchical hidden Markov model that reliably detects rare CNVs overlapping coding exons. GATK-gCNV adjusts for known WES read-depth confounders such as sequence composition and mappability of exon bins, while also adjusting for unspecified technical batch confounders. This algorithm is able to achieve high recall (>90%) from standard WES by comparison to exonic CNVs captured by deep WGS, while also dramatically reducing false positive discoveries to a level appropriate for stringent variant association testing.

To explore the performance of GATK-gCNV in the exome portion of the BGE data, we tested 400 familial samples from an autism cohort, the SSC provided from the SFARI,^9^ all of whom have been independently sequenced with high-coverage genome sequencing, exome sequencing, and have matching microarray data. These samples represented 400 individuals from 100 quartet families, each with two children and both parents, providing a robust design for benchmarking and evaluating *de novo* and transmitted events. We generated BGE data for these samples and applied GATK-gCNV to the exome regions for CNV detection following the published exome CNV parameters.^10^ Using gold-standard CNV calls from high coverage WGS data of these samples, we achieved 87% recall of CNVs spanning 5 or more exons (**Figure 2A**). Recall was slightly lower with BGE compared to standard WES because of lower exome coverage (<60X in BGE vs >60x for previously benchmarked WES). Focusing on only the samples with higher coverage BGE (>60x) achieved 100% recall at 5 exons compared against our truth genomes. GATK-gCNV maintained a ∼90% positive predictive value (PPV) at a resolution of 3 or more exons. Taking advantage of the complete family structure of the data, we also examined the *de novo* CNVs that were detected, and found that 100% of the confirmed *de novo* CNVs (11 *de novos*) reported in the gold-standard WGS CNV study were also detected in BGE data with no additional false positive *de novo* CNVs predicted by our application of GATK-gCNV to BGE data.

**Figure 2.**
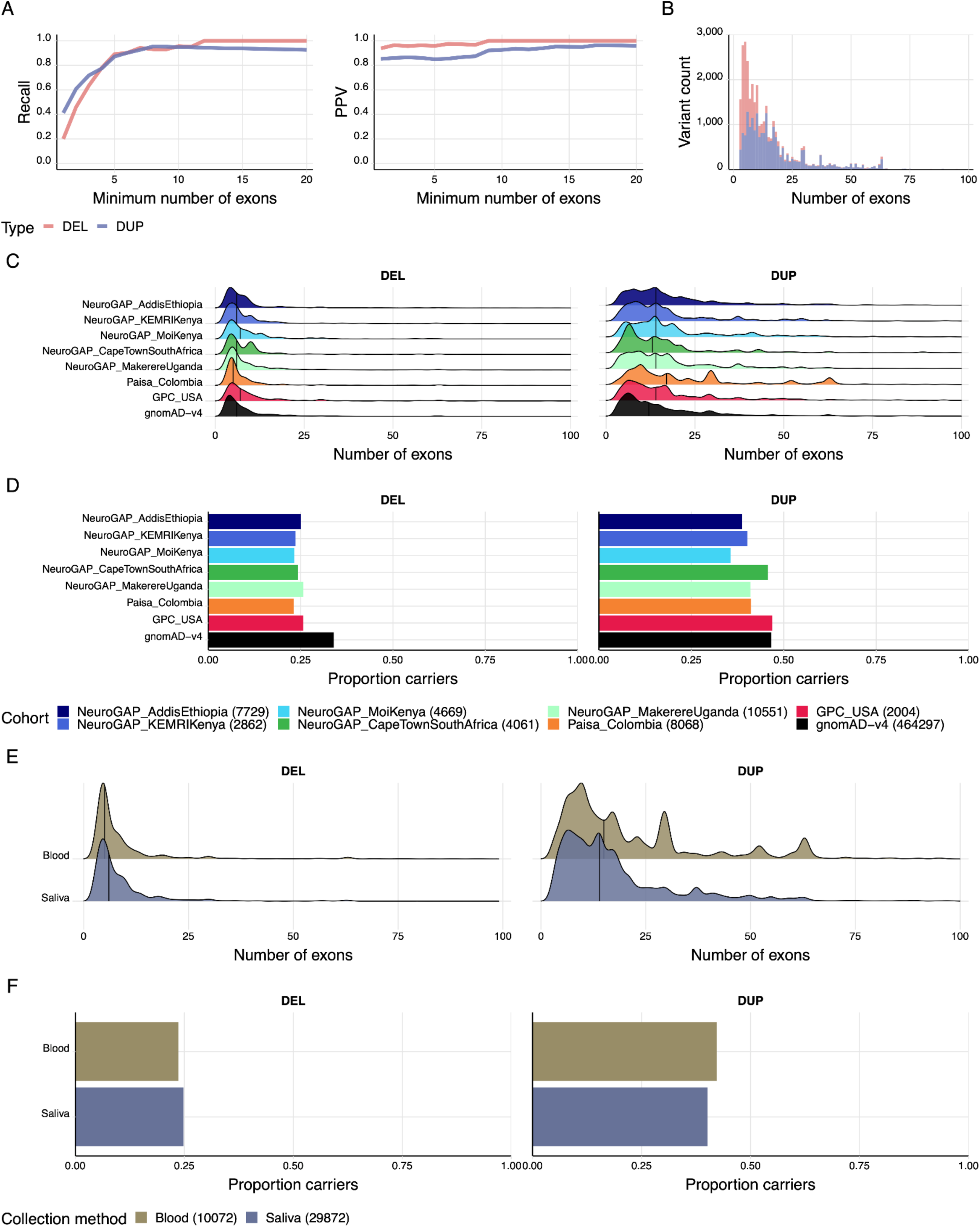
Protein-coding copy number variants have expected qualities with BGE compared to WGS. A) Recall and positive predictive value (PPV) of CNVs called from the BGE relative to matched WGS samples (N=400). B) Distribution of deletions and duplications across all cohorts. C) Distribution of CNV sizes across cohorts by number of exons. D) Proportion of unique deletion and duplication carriers across cohorts. E) Comparison of CNV size across cohorts by number of exons for saliva and blood. F) Comparison of unique deletion and duplication carriers between blood and saliva.

We applied GATK-gCNV to all BGE samples that passed SNV QC. We found that there were more deletions than duplications, and duplications were larger than deletions on average (**Figure 2B**), consistent with previous observations from WGS and WES studies.^11^ Across cohorts, we found similar distributions of CNV size as measured by number of exons covered for both deletions and duplications (**Figure 2C**). Finally, we compared whether CNV discovery with GATK-gCNV differed between blood and saliva, and found no meaningful differences for deletions and duplications in terms of unique carriers or CNV size (**Figure 2E-F**). Similarly, when we restricted to both control samples and constrained genes across two different constraint metrics (LOEUF^12^ and GISMO-mis^13^), we did not find significant evidence for more CNVs in blood than saliva samples (OR = 1.2001, p = 0.1243 for LOEUF; OR = 1.2625, p = 0.0446 for GISMO-mis) (**Supplementary Figure 2**).

### High concordance between imputed BGE and orthogonal GWAS array data generation method

We imputed BGE data directly from CRAM files with GLIMPSE2^14,15^ using a harmonized high-coverage WGS reference panel composed of the Human Genome Diversity Project and 1000 Genomes Project (HGDP+1kGP)^16^. In total, we imputed over 67 million bi-allelic single nucleotide polymorphisms (SNPs) across the allele frequency spectrum. We filtered to high-quality variants, which we define as INFO score ≥ 0.8, resulting in over 30 million SNPs in downstream concordance analyses. The vast majority of the SNPs removed due to INFO-score filtering had minor allele frequency (MAF) < 0.01 (see **Supplementary Table 2**).

Subsets of participants with BGE data from each cohort were also genotyped on the Illumina Global Screening Array (GSA). We evaluated imputation accuracy by comparing BGE imputed genotypes to unimputed GSA array-based genotypes that passed QC (**Methods**), providing an orthogonal comparison of genotypes (**Supplementary Table 3**). We computed Pearson’s aggregated R^2^ within each cohort, grouping SNPs by MAF bin. We found that all cohorts exceed 90% R^2^ across all MAF bins analyzed (**Figure 3**), suggesting high accuracy of the imputed genotype dosages derived from BGE sequencing. We observed a slight downward trend in aggregated R^2^ at MAF> 0.05, though despite this the accuracy remains high.

**Figure 3:**
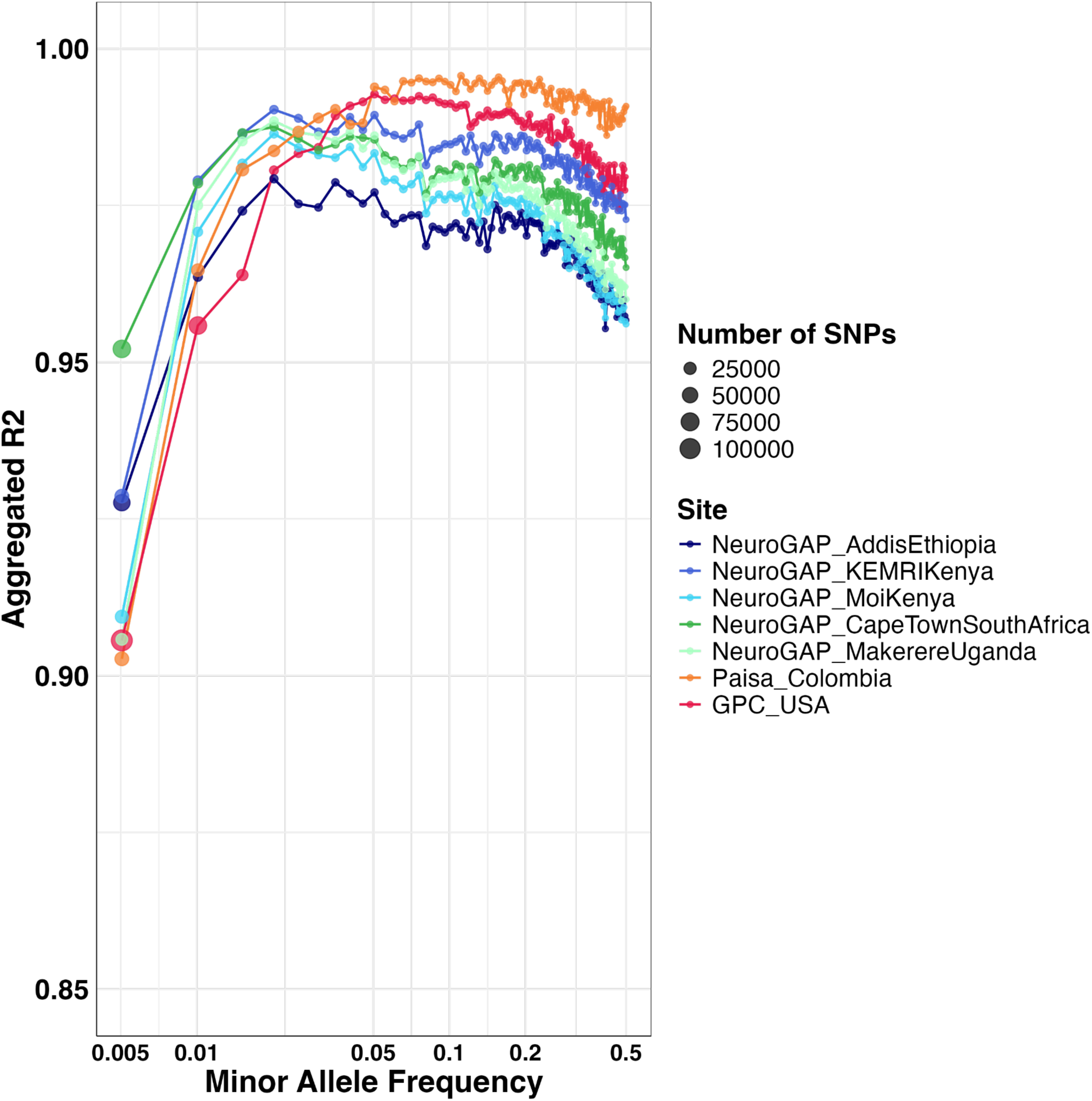
Imputation of BGE data is highly concordant with GWAS array data across MAF bins. The sizes of points correspond to numbers of SNPs in each MAF bin. Variants are filtered to those passing an INFO score >= 0.8. SNP MAFs are defined within cohorts using the GSA array for the Paisa and GPC cohorts. Due to limited GSA samples in the NeuroGAP cohorts, MAFs are defined using the HGDP+1kGP AFR subset.^16^

We also computed non-reference concordance values per cohort (**Supplementary Figure 3**), and found that for common variants (MAF ≥ 1%), BGE data from all cohorts were accurately imputed with at least 92% non-reference concordance. For the lowest MAF bin (0-0.005), we find that all cohorts have a non-reference concordance of at least 83%, suggesting that while there is a slight drop in accuracy, the majority of intermediate to common frequency variants are well-imputed, in agreement with the aggregated R^2^. In addition to INFO score, filtering on posterior genotype probabilities (GP) can improve these concordance metrics (**Supplementary Figures 4A** and **4B**) and attenuate the slight downward trend in aggregated R^2^ with a minimal fraction of dropped genotypes (**Supplementary Figure 4C**). Overall, this analysis shows that imputed genotypes generated from low-pass WGS on the BGE platform are highly concordant with standard array-based genotypes.

### Imputation quality of BGE data across ancestries empowers common variant analysis

We further evaluated BGE imputation accuracy by dividing genotypes into different ancestry tracts based on their diploid local ancestry, following previous work.^17^ We focus on the Paisa and GPC cohort here, as these populations have recently undergone global continental admixture. Using quality controlled GWAS array genotype data, we inferred 3-way local ancestry for Paisa and 2-way local ancestry for GPC using RFMix2^18^. As reference panels for local ancestry inference, we used all individuals within ancestry groups and labels provided by the HGDP+1kGP resource as follows: AFR=African, AMR=Admixed American, and EUR=European populations (AFR/AMR/EUR in 3-way inference and AFR/EUR in 2-way inference). We selected subsets of the HGDP+1kGP^16^ as reference panels for distinct source populations of admixture within Paisa and/or GPC cohorts (see **Methods**).

Aggregate R^2^ stratified by local ancestry backgrounds across MAFs are shown in **Figure 4**. For common variants (MAF ≥ 1%), imputation accuracy is qualitatively similar among various ancestral backgrounds, achieving >90% accuracy in both aggregate R^2^, as well as non-reference concordance (**Supplementary Figure 5A** and **5B**). Aggregated R^2^ and non-reference concordance for heterozygous diploid ancestry genotypes are shown in **Supplementary Figures 5C** and **5D** for the Paisa cohort. For less common SNPs, AMR ancestry has lower R^2^ than EUR and AFR ancestry within the Paisa cohort, a finding consistent with previous local ancestry imputation accuracy measurements.^17^ Similarly, AFR ancestry generally does not perform as well as EUR ancestry within the GPC cohort, as expected. Accuracy differences among rare SNPs are small, with difference in aggregate R^2^ less than 10%. Overall, these results suggest that low-coverage genotypes from the BGE platform can be statistically imputed with high accuracy, even when samples exhibit diverse continental admixture.

**Figure 4.**
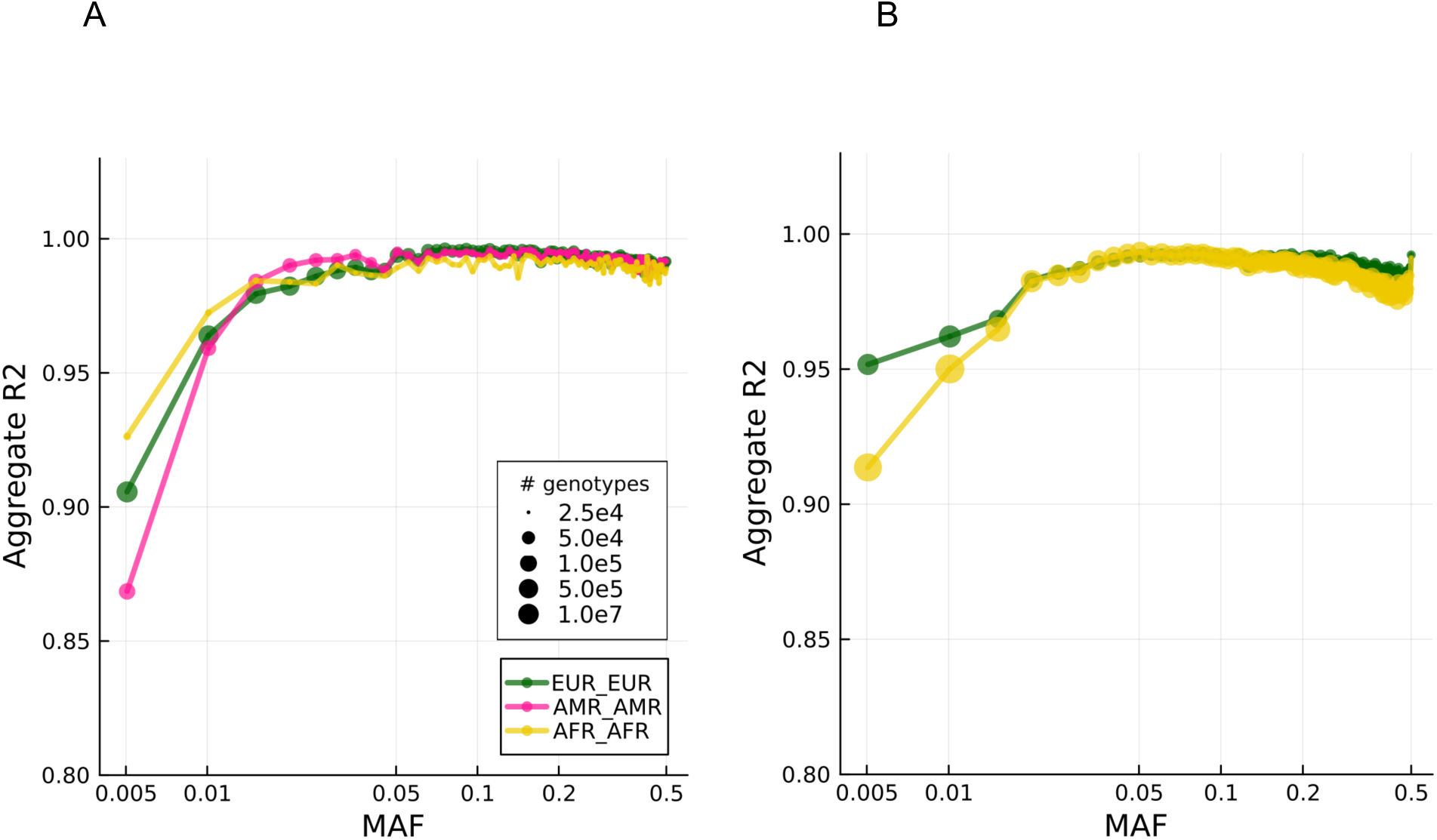
Imputation accuracy differentiates by local ancestry background at low allele frequencies. A) Aggregate R^2^ in the Paisa cohort. B) Aggregate R^2^ in the GPC cohort. Note # genotypes legend stands for number of genotypes, with one value for each sample and each SNP. Heterozygous ancestry results with non-reference concordance measurements are in **Supplementary Figure 5**.

To summarize the balance of cost with variants assayed, we used a subset of NeuroGAP genomes (N=79) with overlapping genetic data from deep WGS, BGE, and Illumina GSA sites to count the number of variants assayed with imputation where relevant in **Table 3** (Methods). We show that BGE captures the vast majority of common (94.4%) and rare (94.7%) coding variants found in 30x WGS, considerably more than can be imputed with GWAS arrays (66.1% and 27.6%, respectively). In the non-coding regions, more comparable fractions were imputed in BGE versus Illumina GSA for common (79.8% vs 67.6%) and rare variants (61.2% vs 24.5%); consistently more variants were imputed with BGE.

**Table 3.**
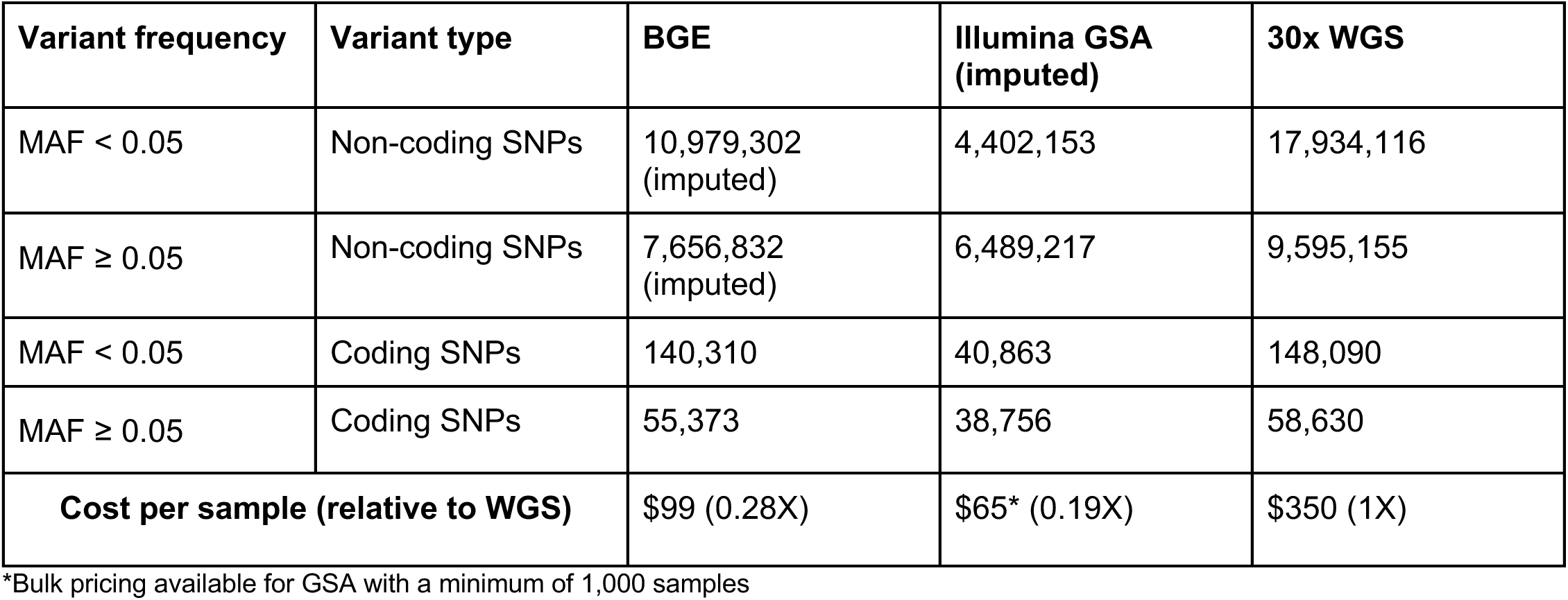
BGE provides a cost-effective balance of variants accurately assayed compared to other data generation strategies. Monomorphic variants were removed, and variants were filtered for those with dosage R^2^ or INFO-score ≥ 0.8 (more details in Methods). Coding and non-coding regions are defined by capture region boundaries for BGE. Costs for BGE and WGS are starting costs described by https://broadclinicallabs.org/ as of August 27, 2024. Notably these costs are basic lab processing costs only that do not include DNA extraction, project management, data storage, analytical, or other costs.

| Variant frequency | Variant type | BGE | Illumina GSA (imputed) | 30x WGS |
| --- | --- | --- | --- | --- |
| MAF < 0.05 | Non-coding SNPs | 10,979,302 (imputed) | 4,402,153 | 17,934,116 |
| MAF ≥ 0.05 | Non-coding SNPs | 7,656,832 (imputed) | 6,489,217 | 9,595,155 |
| MAF < 0.05 | Coding SNPs | 140,310 | 40,863 | 148,090 |
| MAF ≥ 0.05 | Coding SNPs | 55,373 | 38,756 | 58,630 |
| Cost per sample (relative to WGS) | | \$99 (0.28X) | \$65* (0.19X) | \$350 (1X) |

\*Bulk pricing available for GSA with a minimum of 1,000 samples

## Discussion

The BGE sequencing approach represents a significant advance by capturing genetic variation in an unbiased, high-quality, and cost-effective manner. The initial development and testing of BGE was recently described,^8^ and the in-depth analyses here clearly demonstrated that BGE sequencing can be accurately and efficiently scaled to empower human genetics research, especially in diverse and underrepresented populations. By unifying whole genome common variants and coding rare variant analyses from a single sequencing run, our analyses showcase the potential of BGE to enhance genomic discoveries and improve variant detection. In contrast to a recent study which also combined low coverage WGS with high coverage WES,^7^ blending library preparations together at optimized ratios prior to sequencing is unique to the BGE workflow. Our strategy balances variant calling and imputation accuracy at a lower sequencing costs (28% of WGS costs here, more expensive and variable previously). Furthermore, our approach has streamlined workflows, minimizing disjoint sample failures between WGS and WES data, thereby ensuring complete datasets for every successful sample sequenced. We see no clear limitations or biases introduced as a function of blended sequencing, as both deep exome variant identification and low-pass genome imputation show strong concordance when compared against independently sequenced callsets. However, we note that the choice of reference panel when performing imputation of low-pass genome sequencing data can introduce biases with respect to imputation accuracy. For example, we found lower aggregate R^2^ in AMR haplotypes from the Paisa population due to fewer Amerindigenous haplotypes in admixed AMR individuals relative to EUR and AFR ancestry individuals included in the HGDP+1kGP reference panel. Relatedly, while imputation panels that support GWAS arrays have increased in scale and diversity far beyond HGDP+1kGP, low-coverage imputation requires individual-level haplotype data and uses methods not currently supported by existing servers.^14^

The BGE technology delivers consistent coverage and quality across cohorts, with high accuracy in calling coding CNVs and only minor differences in coverage based on saliva versus blood-based collection. In future saliva-based BGE sequencing, the protocol can be altered to draw a slightly larger aliquot in saliva samples to overcome this discrepancy. We find high recall and PPV when calling coding CNVs, with no discernable impact of saliva versus blood collection on CNV call rates. The slightly lower recall in calling coding CNVs overlapping < 5 exons is due to the lower BGE target deliverables (90% of exome target reaching 10x depth) compared to deep exome protocols performed at the Broad Institute (85% of exome target reaching 20x depth). These benchmarks of comparability between protein-coding CNVs with BGE versus WES demonstrate a unique strength of the technology at this price point regarding assayable variants. However, additional work is underway to further quantify the full breadth of genome-wide SV detectable with low-pass WGS read data from BGE. While segregating SVs can be tagged to some extent by SNPs^11^, the lack of an SV imputation service limits widespread adoption, and most rare and *de novo* SVs would never be detected with imputation.

The BGE technology excels in common variant imputation from low-pass WGS reads, demonstrating consistently high concordance with GWAS array data across diverse cohorts, and within specific local ancestry tracts from large recently admixed cohorts. As low-pass WGS data is not biased towards a specific set of selected variants, the performance should remain robust and flexible compared to GWAS arrays across all varieties of future imputation reference panels, making it a powerful long-term tool for common genetic variant association studies. While GWAS array data is the most cost-comparable alternative to BGE, we acknowledge that it is not a gold standard for benchmarking imputation accuracy due to its limited coverage and occasional errors.^19^ We aimed to retain only the highest quality genotypes in the selected array SNPs, but discrepancies arising between BGE and array data cannot be attributed solely to noise from low-pass imputation of BGE data. BGE is a particularly appealing genetic data strategy because it has similar costs to a GWAS array with genomic coverage akin to an exome plus array with imputation; deep WGS data, a gold standard, only finds ∼1% more single-variant and gene-based associations than an exome plus array despite costing >3.5x more than BGE.^20^

Our evaluation across a highly ancestrally diverse suite of cohorts establishes BGE sequencing as a technology that bridges the gap between cost, comprehensive coverage, and data quality for large-scale genomic studies. Its high-quality exome data, reliable CNV calling, and accurate whole-genome imputation across diverse populations make it a valuable tool for advancing genomic research. By enabling more inclusive studies, BGE technology has the potential to enhance our understanding of genetic variation and its implications for human health around the world.

## Methods

### Wet lab protocol

#### Biological Samples

To date, BGE DNA samples have been extracted from saliva and whole blood specimens outside of our lab. Samples derived from whole blood are generally preferred. Lower alignment rates of sequencing reads have been observed in saliva samples due to the presence of bacterial species. Furthermore, saliva samples can be more difficult to accurately quantify due to their consistency and heterogeneity. Following DNA extraction, we follow the BGE protocol described previously and detailed briefly below.^8^

#### PCR-Free Library Preparation

After an initial quantification, DNA is normalized to 50 ng/uL and transferred into a 384-well plate. Normalized DNA is purified using an automated 2.75X solid phase reversible immobilization (SPRI) clean-up with Ampure XP Agencourt beads (Beckman Coulter, Indianapolis, IN, USA). Cleaned DNA is then quantified by spectrophotometry (Lunatic, Unchained Labs, Pleasanton, CA, USA) and normalized to 25 ng/uL. DNA (target of 134 ng input) undergoes a reduced and customized fragmentation/end-repair/A-tailing reaction for Illumina-compatible PCR-free library construction with custom NEBNext Ultra II FS DNA Library Preparation Kits (New England Biolabs) using the following conditions (37℃ for 42.57 min, 65℃ for 30 min). Unique, dual-indexed adaptors (NEBNext Unique Dual Index UMI Adaptors, New England Biolabs) are ligated to fragments (20℃ for 20 min) and libraries undergo two consecutive SPRI size-selections (0.5x and 0.55x). Cleaned and size-selected PCR-free libraries are quantified by qPCR (Kapa Library Quantification Kit, Roche), then normalized (to ∼0.3-2 nM, depending on concentrations) before pooling into a single tube and concentrated (Unagi, Unchained Labs, Pleasanton, CA, USA).

#### Exome Captured Library Preparation

An aliquot from the pre-normalized and pre-pooled PCR free libraries is used as input for PCR amplification (cycles are 98℃ for 30 sec, 12 cycles of [98℃ for 10 sec, 65℃ for 75 sec], 65℃ for 5 min) using the NEBNext Ultra II FS Library Preparation Kit and primers from the indexed adaptor kit (New England Biolabs). PCR amplified libraries are quantified by spectrophotometer (Lunatic), purified with a 1X SPRI-cleanup (Ampure), and normalized to 70 ng/uL. Samples are then pooled and undergo exome capture (Twist Alliance Clinical Research Exome probes from Twist Biosciences, South San Francisco, CA, USA) using the recommended hybridization-capture protocol for xGen Hybridization Capture Core Reagents (Integrated DNA Technologies, Coralville, IA, USA).

#### Blending of PCR-Free Genome and Exome Captured Libraries

PCR-free and exome-captured pools are both qPCR-quantified on the same qPCR run. Nanomolar concentrations are taken into account to calculate the appropriate volumes to blend 33% WES with 67%WGS. BGE samples are again qPCR quantified for sequencer loading calculations.

#### Sequencing

BGE blended pools with 384 samples containing unique barcodes are sequenced across 6 lanes of NovaSeq S4 (Illumina) with 2x150 bp runs.

### Datasets analyzed to evaluate BGE quality

BGE data used in these analyses were generated at the Broad Clinical Lab. Sample cohorts included in the dataset were recruited and submitted from the collaborating institutions for the PUMAS project which includes cohorts from the GPC, Paisa population, and NeuroGAP-Psychosis, as described below, in **Table 2**, and in **Supplementary Table 1**.

The Genomic Psychiatry Cohort (GPC) is a multi-institutional collaboration led by Rutgers University. The GPC resource includes an NIMH-managed repository of genomic samples at SAMPLED, genotypic and sequence data, and detailed clinical and demographic data for investigations of schizophrenia, bipolar disorder, OCD, and COVID from a variety of ancestries collected in the USA. Within this analysis, we included 4,553 samples with 3,926 passing QC filters.

The Paisa population is a genetic isolate from Colombia that has expanded rapidly following a series of migration-related bottlenecks. They have been the focus of genetics studies in neuropsychiatric disorders in the last decade. Paisa BGE data generated here emerged from a long-standing collaboration between teams at the Universidad de Antioquia, Medellin, Colombia and University of California, Los Angeles to recruit samples from this population. Within the analysis, we included 9,007 samples with 8,200 passing QC filters.

The Stanley Center at the Broad Institute of MIT & Harvard initiated the NeuroGAP-Psychosis project in 2015, a collaboration with colleagues at Addis Ababa University in Ethiopia, KEMRI-Wellcome Trust in Kenya, Makerere University in Uganda, Moi University/Moi Teaching and Referral Hospital in Kenya, the University of Cape Town in South Africa, and the Harvard T.H. Chan School of Public Health in the United States. With initial plans to recruit 35,000 participants (half cases with a diagnosis of schizophrenia or bipolar disorder and half controls), the target was expanded to 39,000 participants via the PUMAS Project awarded by NIMH. Sample recruitment ultimately exceeded the revised target, with >42,000 samples collected across the five NeuroGAP-Psychosis collection sites. Within the analysis, we included 39,886 samples with 35,138 passing QC filters.

Data from the PUMAS project are being deposited at the NIMH Data Archive (NDA). At the time of manuscript submission, genomic data from the first 10,000 samples have been submitted. All remaining samples from the PUMAS grant will be submitted at the end of the grant period. Data that are designated with NDA-GRU data use will be deposited into DNA collection #3805. Data designated with Disease-Specific (Mental Health) - DS (Mental Health) will be deposited into DNA collection #4538. Finally, data designated Health/Medical/Biomedical, NDA-HMB-MDS will be deposited into DNA collection #4539.

### Exome Quality Filtering

We conducted quality control filtering to restrict to sites and variants with high confidence in exome data. We filtered out sites with >6 alleles that failed VQSR, were within low-complexity regions, or outside of Twist target capture regions. Within an individual, genotype calls were filtered if the read depth <10X, genotype quality <20, or if allele balance was <0.2 or >0.8 in heterozygous calls, or if allele balance was <0.8 in homozygous alternate calls.

We filtered samples based on WES and WGS coverage, ancestry, and WES sample quality metrics to restrict to high-quality samples for subsequent analysis. First, we removed samples with low WES or WGS coverage. Exome coverage per sample was determined by calculating the fraction of the exome target covered with at least 10x read depth. Samples with exome fractions less than 90% or estimated WGS coverage of less than 1x were removed. Additionally, samples with chimeric or contamination read rates greater than 5% were removed, resulting in 853 removed samples.

Next, we determined genetic ancestry of the PUMAS samples using the quality filtered WES portion of the BGE. We combined the PUMAS data with four reference panels with diverse ancestries: HGDP, 1KGP, AWI-GEN, and the African Genome Variation Project (AGVP).^16,21,22^ Throughout this manuscript, we use ancestry labels assigned by existing genomic reference panels, including: EUR (European), AFR (African), and AMR (Admixed American - an imprecise label introduced by the 1000 Genomes Project to describe individuals with recent admixture from multiple continents including Amerindigenous ancestry). Sites overlapping between the PUMAS data and all reference datasets were filtered to biallelic variants with call rates >0.98 and minor allele frequencies >0.1%. We conducted LD pruning to extract independent markers and calculated the top 10 principal components across all samples using Hail’s hwe_normalized_pca function. We fit a random forest algorithm using the top 10 principal components to the reference datasets of known ancestry and then applied the algorithm to the PUMAS data. We required PUMAS samples to have a probability of at least 0.7 to assign ancestry. Samples with probabilities less than 0.7 were removed (N removed = 1,799).

We calculated sample quality metrics using Hail’s sample_qc function. Within each ancestry and cohort combination, we removed samples with outlier values defined as values >4 median absolute deviations from the mean for the following metrics: N singletons, N insertions, N deletions, N transitions, N transversions, heterozygosity ratio, transition to transversion ratio, and insertion to deletion ratio. Sample quality metrics filtering removed 2,317 samples.

Finally, we removed samples with discrepancies between imputed genetic sex and reported gender. Genetic sex was imputed separately for each genetic ancestry group by calculating the inbreeding coefficient on the X chromosome using common, independent markers after removing markers within the pseudoautosomal region. Samples with an inbreeding coefficient <0.6 were labeled ‘female’ and samples with an inbreeding coefficient >0.6 were called ‘male’. We removed samples whose imputed sex and reported gender did not match, resulting in 316 removed samples.

### Copy number variant calling and quality control

#### CNV calling

To call copy number variants from the deep coverage exome data, GATK-gCNV was used on the exome intervals following a previously described pipeline.^10^ GATK-gCNV adjusts for known WES read-depth confounders such as GC content and mappability, while simultaneously adjusting for unspecified technical batch confounders such as sample extraction, sequencing, library preparation, and mapping quality. We took the read-depth data as input from the BGE samples over a set of canonical transcript genomic intervals. Briefly, the raw sequencing files were compressed into counts of the reads over the set of annotated exons and used as input into GATK-gCNV. A principal component analysis (PCA)-based approach was used on the compressed observed counts to identify differences in the capture kit. Within each cohort, PCA was used to define ancestrally similar batches of 1,000 samples; from each, a random subset of 200 samples were used to train a CNV-discovery model tailored to each batch. Subsequently, a distance-based and hybrid-density-based clustering approach was used to curate samples into batches to process in parallel. Once batching determination was completed, GATK-GCNV was used on each batch and metrics for filtering were produced by the Bayesian model underlying the data to balance recall and positive predictive value.

#### CNV benchmarking

To benchmark BGE CNV exome data, we used data from 100 quads from the Simons Simplex Collection (SSC)^23–25^, sequenced previously. Benchmarking was carried out on all rare CNVs (frequency <1%). The 100 quads have existing CNV calls^24,25^ from deep whole-genome sequencing (∼30X), and the CNV calls from WGS were used as a gold standard to benchmark the BGE CNVs. Recall was calculated by the proportion of WGS data sites that had a match in the BGE CNV callset. Moreover, for any given site, if at least 50% of samples that had that variant in the WGS data also had a GATK-gCNV call with consistent directionality (duplication or deletion) that overlapped at least 50% of captured intervals, this was considered a correct call. The optimal recall and PPV was for CNVs >4 exons in the 100 quads. Samples were removed if there were >10 rare CNVs, defined as <1% frequency across the cohort. Any samples with >10 high quality CNVs were removed. We additionally removed samples that failed SNV quality control. Subsequently, CNVs were filtered to have a QS (Quality Score) >200 and restricted to <1% frequency for each ancestry. We further benchmarked the performance of *de novo* CNVs derived from BGE data using GATK-gCNV as a function of the number of captured exons of canonical transcripts compared to validated WGS *de novo* CNVs from the quads. We additionally benchmarked the CNVs in saliva and blood against two independent constrained gene sets (top 1000) based on LOEUF^12^ from gnomAD v2 and GISMO-mis.^13^

#### Imputation

To impute the low-coverage whole genome sequencing data, we used the Genotype Likelihoods IMputation and Phasing (GLIMPSE2) method.^14^ We used the HGDP+1kGP reference panel^16^ after filtering out singleton variants and indels, resulting in about over 67 million variants available to be imputed.

To phase and impute this large-scale data in a cost-effective manner, we used Broad’s Hail Batch Service to submit randomized batches of 200 individuals. Then, we merged imputed results from the batches using bcftools. In total, we imputed over 47k samples for just over $17,000, averaging about $0.36 per sample (**Supplementary Table 4**).

Allele frequencies (AF) were corrected after merging by calculating a weighted mean of the batch AFs based on individual batch values (for *i* batches and *N* samples per batch):

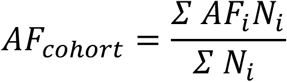

Similarly, INFO scores (*I*) were corrected post-merge via:

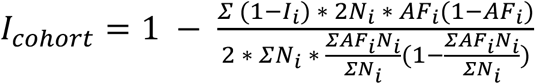

This process resulted in a final set of chromosome-specific BCF files, with all individuals of a cohort included and allele frequencies and INFO scores updated to reflect the full sample size.

### Quality Control of GSA data

Variants genotyped via the Illumina Global Screening Array (GSA) platform were first filtered for a SNP call rate of at least 95%, then samples were filtered to have a call rate of at least 98%, heterogeneity F statistic or inbreeding coefficient |F_HET_| < 0.20, and concordance between genetically inferred and reported sex. A second SNP filtering step with a call rate of at least 98% was performed after the sample quality control, followed by filtering steps to remove SNPs with missingness differences > 2% between cases and controls, and Hardy-Weinberg equilibrium (HWE) p-value of 1e-06 for controls and 1e-10 for cases. We used King^26^ to calculate the genetic relationship matrix and removed one sample in any pairs of individuals with second-degree or closer relatives. Finally, we used the conform-gt tool provided by the Beagle imputation software^27^ to align the strand and allele order within the GSA VCFs to the HGDP+1kGP reference panel.^16^

### Concordance between BGE and GSA GWAS array data

To evaluate BGE data accuracy post-imputation, we compared BGE with previously generated Illumina GSA data as ground truth on the subset of individuals with both data types (**Supplementary Table 3**). Imputed variants were filtered to match those in the GSA. We computed non-reference concordance and aggregated R^2^ as a function of MAF for each cohort. To evaluate non-reference concordance^28^, we define it as the number of true positives divided by the sum of the true positives, false negatives, and false positives (excluding missing sites and SNPs with only reference alleles within the GSA subset), as shown in the formula below. We computed aggregated R^2^ per MAF bin by stacking all SNP dosages (from the imputed data) and genotype calls (from the GSA data) within a minor allele frequency bin, and calculating the squared Pearson’s correlation coefficient. Note that to avoid discrepancies in minor allele frequency for the NeuroGAP subsets (each with around 160 individuals), minor allele frequency values were based on the HGDP+1kGP AFR subset rather than in-sample allele frequencies. For the larger datasets, allele frequency values were taken from in-sample calculations.

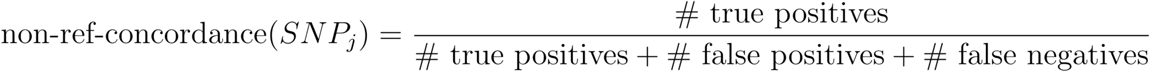

Let *X_GSA_* be the matrix of genotypes for a given set of individuals and SNPs based on GSA array data, while *X_BGE, imputed_* is the matrix of imputed dosages for the same set of SNPs and individuals. We can calculate aggregated R^2^ per a given MAF bin for a SNP as follows^29^:

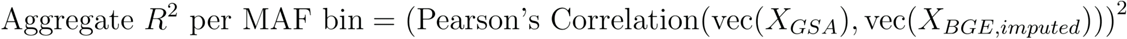

### Concordance between BGE and GSA by Local Ancestry Inference (LAI)

We also evaluated post-imputation accuracy in BGE data stratified by each genotype’s local ancestry background. We focus specifically on the Paisa and GPC populations, which have seen recent admixture between European, Native American/Amerindigenous, and African continental ancestry. We first phased array genotypes via Beagle 5.4^30^ using the full HGDP+1kGP reference panel (n=4099).^16^ Next, we followed the Tractor^31^ tutorial and computed local ancestry with RFMix2 software.^18^ For the Paisa cohort, we modeled 3-way local ancestry to capture admixture represented by European (EUR), African (AFR), and Admixed American (AMR, as a proxy for Amerindigenous ancestry) populations in the HGDP+1kGP reference panel.

Specifically, we filtered the reference panel to only samples with AFR/EUR/AMR population labels. For LAI to be most useful, reference samples themselves should be representative of ancestral populations and non-admixed. Because AMR samples tend to themselves be recently admixed, we excluded 358/549 reference AMR samples that have <90% Amerindigenous ancestry as measured by RFMix2. As a result, the total number of reference samples N_total_=1935 are split among 191 AMR, 752 EUR, and 992 AFR. For the GPC population, we inferred 2-way local ancestry to capture European and African admixture. When running RFMix2, we added the optional EM flag set to run 1 iteration to account for remaining admixture within the subsetted reference panel. Finally, we also included the ‘-n 5’ flag to account for reference panel sample size imbalance.

After LAI, two metrics were used to evaluate imputation accuracy for different ancestry backgrounds, including aggregate R^2^ and aggregate non-reference concordance, computed by stacking all SNPs within a MAF bin and evaluating a single non-ref concordance value. This is analogous to the aggregate R^2^ formula except we compute concordance rather than squared correlation after stacking SNPs column-wise. Note that we favor the *aggregate* non-reference concordance metric as opposed to averaging SNP-by-SNP concordances due to the presence of rare SNPs. This is because a rare SNP may have only several copies of the minor allele, and by chance, they may all reside on one specific ancestry background. This leaves no copies of minor alleles for other ancestry backgrounds. Averaging SNP-by-SNP non-reference concordances will therefore include excess zeros for the remaining ancestries, artificially deflating their accuracy. Stacking rare SNPs rescues concordance for rare SNPs among all ancestry backgrounds by virtue of including more copies of the minor allele, ultimately producing more accurate and interpretable results.

Scripts used for this computation can be found here: https://github.com/atgu/bge_analysis/tree/main/concordance/julia#aggregate-r2-based-on-different-local-ancestry-backgrounds

### Number of variants per data generation type

We evaluated a subset of samples from the NeuroGAP data (N=79) that were assayed across three different data generation strategies, including BGE, Illumina GSA arrays, and 30X WGS. To count the number of variants per MAF category in the noncoding BGE data, we used the data imputed with GLIMPSE2 after filtering for an INFO score of at least 0.8 and removing monomorphic variants. For the coding regions, we utilized the high coverage exome data rather than the imputed data, filtering for VQSR PASS variants as well as excluding locus control regions (LCRs) and filtering for monomorphic variants. We required genotypes to have genotype quality (GQ) > 50 and depth > 13, and normalized Phred-scaled likelihoods (PL) to be > 50. These same filters were applied for both coding and noncoding regions in the WGS dataset. For the Illumina GSA GWAS array data, we used the imputed data generated previously^3^ using the phase 3 1000 Genomes Project reference panel with BEAGLE v5.1 for all categories, filtering for dosage R^2^ of at least 0.8 and removing monomorphic variants. Coding regions were defined by Twist target capture region intervals.

## Software availability

Code for conducting imputation and concordance analyses are available at https://github.com/atgu/bge_analysis

Code for running the gCNV pipeline is available at: https://github.com/broadinstitute/gatk/tree/4.1.0.0/scripts/cnv_wdl/germline

## Author contributions

**Imputation Analysis:** T.A. Boltz, B.B. Chu, L. Majara, S. Rubinacci, M. Gatzen, C. Kachulis, S. Kim, L.L. Nkambule, S.J. Parsa, E. Stricker, M.T. Yohannes. **CNV Analysis:** C. Liao, R. Ye, J.M. Fu. **SV Analysis:** L. Zhan, S. Lee, S. Mangul. **QC Analysis:** C. Liao, J.M. Sealock, R. Ye, S.E. Medland. **Data Generation:** A.B. Bradway, M.L. Gildea, T.C. Hill, K.M. Hubbard, P.R. Kalmbach, C.R. O’Neil, A.M. Olivares, F.L. Reagan, J.A. Schneider, J. Tang. **Data Contributors:** T. Abebe, D. Akena, M. Alemayehu, F.K. Ashaba, L. Atwoli, A. Fekadu, S. Gichuru, W.E. Injera, R. James, M. Joloba, R. Kamulegeya, S.M. Kariuki, G. Kigen, N. Koen, E.K. Kwobah, J. Kyebuzibwa, J. Makale, J. McMahon, P. Mowlem, H. Musinguzi, R.M. Mwema, N. Nakasujja, C.P. Newman, C.R.J.C. Newton, C.M. Olsen, L. Ongeri, A. Pretorius, R. Ramesar, W. Shiferaw, D.J. Stein, A. Stevenson, R.E. Stroud II, S. Teferra, D. Whiteman, Z. Zingela. **Program Management:** S.B. Chapman. **Wet Lab Leadership:** M. DeFelice, J.L. Grimsby, N.J. Lennon. **Analytical Supervision:** E.G. Atkinson, T. Bigdeli, H. Brand, L.B. Chibnik, D.P. Howrigan, H. Huang, K.C. Koenen, E.A. Lopera-Maya, C.N. Pato, M.T. Pato, C. Sabatti, M.E. Talkowski, C. Service, R.A. Ophoff, L.M. Olde Loohuis, N. Freimer, M. Yu, K. Yuan. **Analytical Supervision & Data Contributors:** B. Gelaye, K.C. Koenen, C. Lopez-Jaramillo, R.A. Ophoff, L.M. Olde Loohuis. **Analytic Supervision on Pilot Data:** H. Huang. **Analysis of Pilot Data during Development:** M. Yu, K. Yuan. **Leadership:** B.M. Neale, A.R. Martin.

## Acknowledgements

The blended genome-exome data in this manuscript were generated at the Broad Institute’s Genomics Platform (now Broad Clinical Labs) with support from the National Institute of Mental Health (NIMH) under the Populations Underrepresented by Mental illness Association Studies (PUMAS) grant U01MH125047 to the Broad Institute of MIT and Harvard. This work is supported by a Collaborative U01 grant from the National Institute of Mental Health: *Powering Genetic Discovery for Severe Mental Illness in Latin American and African Ancestries* awarded to: The Broad Institute of MIT and Harvard (U01MH125047); Harvard T.H. Chan School of Public Health (U01MH125045); University of California Los Angeles (U01MH1250452); and Rutgers University (U01MH125049). Structural variant methods development was also supported by NIMH R01115957 and NICHD R01HD081256.

Sample recruitment and analysis of the Paisa_Colombia study was supported through NIMH R01MH113078. Sample recruitment and analysis in the NeuroGAP-Psychosis study was provided by The Stanley Family Foundation as well as R01MH120642.

This research was funded in part by BD^2^: Breakthrough Discoveries for thriving with Bipolar Disorder. For the purpose of open access, the author has applied a CC BY 4.0 public copyright license to all Author Accepted Manuscripts arising from this submission.

C.L. is funded by the Canadian Institutes of Health Research Banting Fellowship. A.R.M. is supported by K99/R00MH117229. E.G.A. is supported by R01HG012869. J.M.F. is supported by NIMH K01MH137407 and Mass General Neuroscience Transformative Scholar Award.

