## Supplementary Tables and Figures for "A blended genome and exome sequencing method captures genetic variation in an unbiased, high-quality, and cost-effective manner"

**Supplementary Table 1.** Cohort breakdown of data included in these analyses with phenotype and recruitment information.

The first institution listed in the “Submitted by” column was responsible for recruiting participants; the second institution was the US-based NIH prime institution. Phenotypes are: SCZ=schizophrenia, BP=bipolar, MDD=major depressive disorder. # Psychosis cases refer to instances that do not fit other diagnoses. Quality control includes filtering on WES and WGS coverage, genetic ancestry, outlier filtering on sample quality metrics, and checking for discrepancies between genetic sex and reported gender.

| Cohort name | Country of Origin | Tissue | Submitted by | # Total Pre-QC | # SCZ Cases | # BP Cases | # MDD Cases | # Psychosis Cases | # Controls | # Total Post-QC |
|---|---|---|---|---|---|---|---|---|---|---|
| NeuroGAP - Addis Ababa University | Ethiopia | Saliva | AAU/ HSPH | 11,715 | 4,171 | 1,209 | 0 | 16 | 5,631 | 11,027 |
| NeuroGAP - KEMRI | Kenya | Saliva | KEMRI/ HSPH | 3,078 | 729 | 485 | 0 | 407 | 1,268 | 2,889 |
| NeuroGAP - Moi Teaching and Referral Hospital | Kenya | Saliva | Moi/ HSPH | 5,040 | 1,385 | 968 | 0 | 2 | 2,361 | 4,716 |
| NeuroGAP - University of Cape Town | South Africa | Saliva | UCT/ HSPH | 8,747 | 2,015 | 710 | 0 | 34 | 3,020 | 5,779 |
| NeuroGAP - Makerere University | Uganda | Saliva | Makerere Uni/ HSPH | 11,306 | 1,501 | 2,784 | 1 | 955 | 5,485 | 10,727 |
| Paisa | Colombia | Blood | UdeA/ UCLA | 9,007 | 1,369 | 2,984 | 2,539 | 22 | 1,284 | 8,200 |
| Genomic Psychiatry Cohort (GPC) | USA | Blood | Rutgers | 4,553 | 906 | 865 | 30 | 2 | 1,865 | 3,926 |
| | | | **Total** | **53,446** | 12,076 | 10,005 | 2,570 | 1,438 | 20,914 | **47,264** |

**Supplementary Table 2.** Number of imputed SNPs by minor allele frequency.

| MAF bin | NeuroGAP unfiltered | NeuroGAP INFO>=0.8 | GPC unfiltered | GPC INFO>=0.8 | Paisa unfiltered | Paisa INFO>=0.8 |
|---|---|---|---|---|---|---|
| **[0,0.01]** | 52,969,838 | 15,808,286 | 53,368,842 | 24,003,218 | 58,363,478 | 23,364,001 |
| **(0.01,0.02]** | 3,381,317 | 3,337,758 | 2,986,989 | 2,972,371 | 1,775,145 | 1,766,482 |
| **(0.02,0.03]** | 1,681,557 | 1,670,725 | 1,599,805 | 1,594,889 | 733,280 | 731,040 |
| **(0.03,0.04]** | 1,084,382 | 1,079,477 | 1,069,749 | 1,067,513 | 452,302 | 451,298 |
| **(0.04,0.05]** | 790,740 | 788,116 | 793,965 | 792,804 | 352,733 | 352,164 |
| **(0.05,0.1]** | 2,249,890 | 2,244,708 | 2,319,863 | 2,318,137 | 1,177,405 | 1,176,502 |
| **(0.1,0.15]** | 1,247,955 | 1,246,018 | 1,271,570 | 1,271,093 | 816,420 | 816,179 |
| **(0.15,0.2]** | 876,932 | 875,664 | 890,479 | 890,207 | 669,883 | 669,746 |
| **(0.2,0.25]** | 667,141 | 666,200 | 669,093 | 668,921 | 572,871 | 572,800 |
| **(0.25,0.3]** | 548,769 | 548,037 | 539,499 | 539,384 | 514,843 | 514,792 |
| **(0.3,0.35]** | 480,545 | 479,879 | 475,999 | 475,875 | 472,541 | 472,504 |
| **(0.35,0.4]** | 433,172 | 432,526 | 431,230 | 431,144 | 458,325 | 458,288 |
| **(0.4,0.45]** | 406,113 | 405,543 | 403,343 | 403,235 | 434,695 | 434,664 |
| **(0.45,0.5]** | 398,198 | 397,559 | 396,123 | 396,019 | 422,628 | 422,584 |
| **Total** | 67,216,549 | 29,980,496 | 67,216,549 | 37,824,810 | 67,216,549 | 32,203,044 |

**Supplementary Table 3.** Number of overlapping samples per cohort with BGE and GSA array data.

| Dataset | N samples with BGE and GSA data |
|---|---|
| NeuroGAP - AddisEthiopia | 158 |
| NeuroGAP - KEMRIKenya | 183 |
| NeuroGAP - MoiKenya | 157 |
| NeuroGAP - CapeTown South Africa | 162 |
| NeuroGAP - Makerere Uganda | 178 |
| Paisa - Colombia | 1,191 |
| GPC - USA | 3,932 |

**Supplementary Table 4.** Costs of imputation per cohort.

| Cohort | Sample Size | Cost | Cost Per Sample |
|---|---|---|---|
| **NeuroGAP** | 35,279 | $12,763.23 | 0.361 |
| **Paisa** | 8,317 | $3,025.45 | 0.363 |
| **GPC** | 3,967 | $1,406.46 | 0.354 |
| **Total** | **47,563** | **$17,195.14** | **0.361** |

### Supplementary Figures

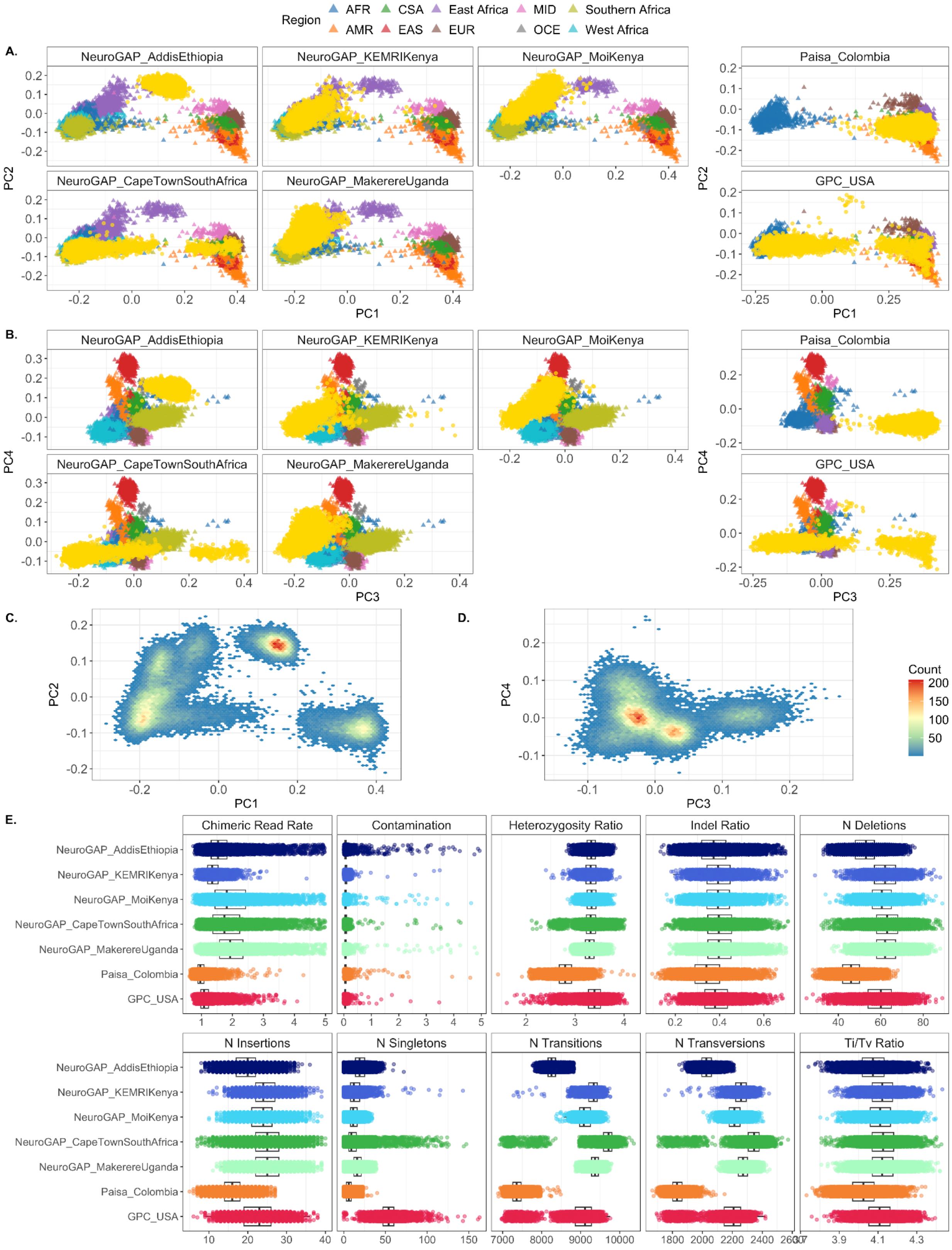

**Supplementary Figure 1.** Principal components with reference panels for all PUMAS cohorts and additional sample quality control metrics.

A. PC1 vs PC2 for PUMAS samples (yellow) and reference samples stratified by cohort. For NeurGAP sites, reference samples are color-coded by region, including four reference panels with diverse ancestries: HGDP, 1KGP, AWI-GEN, and AGVP. For Paisa and GPC, reference panels were limited to HGDP and 1KGP.
B. PC2 vs PC3 for PUMAS samples (yellow) and reference samples stratified by cohort. For NeurGAP sites, reference samples are color-coded by region, including four reference panels with diverse ancestries: HGDP, 1KGP, AWI-GEN, and AGVP. For Paisa and GPC, reference panels were limited to HGDP and 1KGP.
C. Density plots of PC1 vs PC2 for all PUMAS samples.
D. Density plots of PC3 vs PC4 for all PUMAS samples.
E. Estimated chimeric read rates, contamination rates, heterozygosity ratio, indel ratio, number of deletions and insertions, number of singletons, number of transitions and transversions, and Ti/Tv (transition/transversion) ratio per sample per cohort.

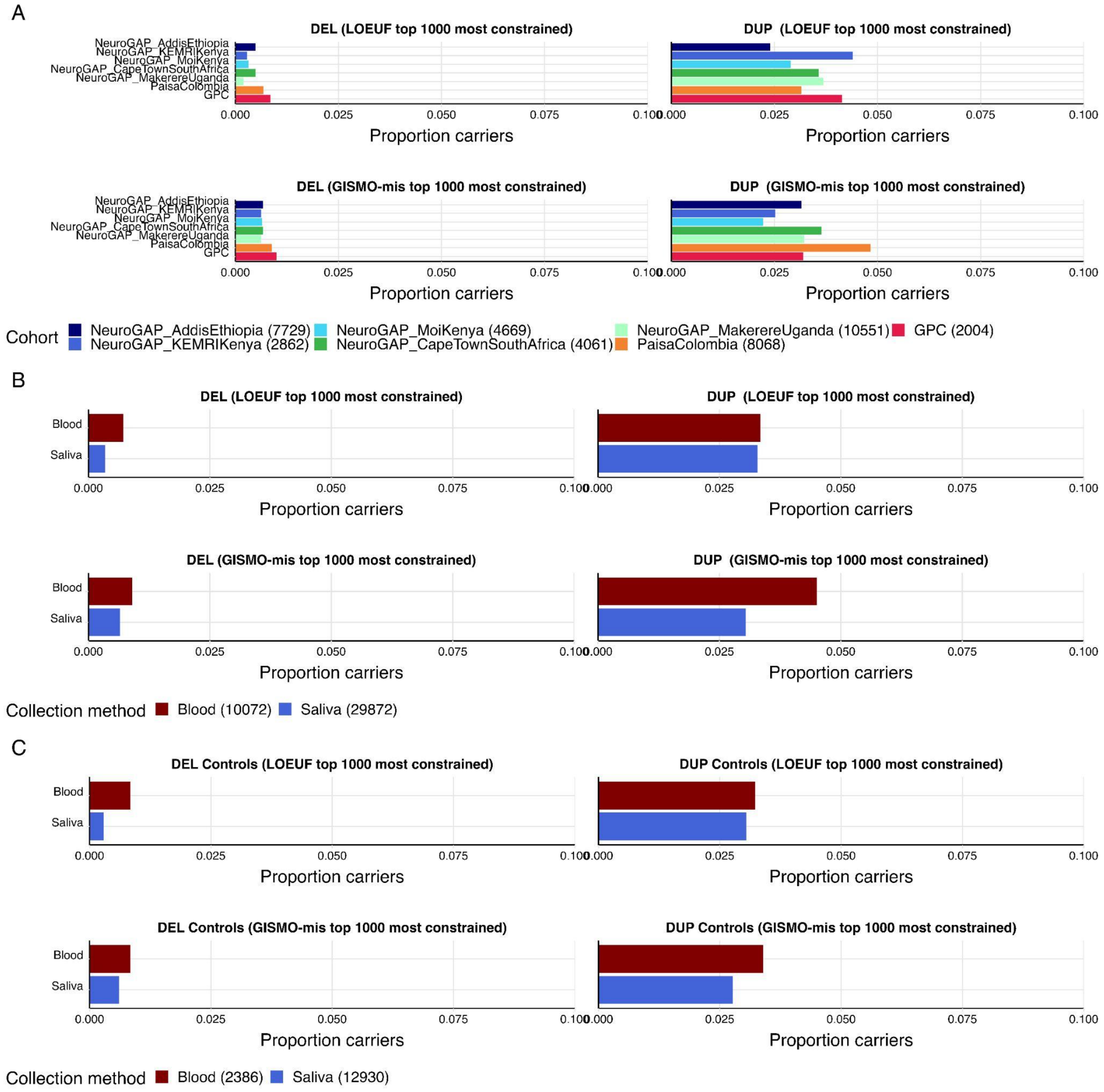


**Supplementary Figure 2.** Proportion of CNV carriers across the top 1000 most constrained genes for LOEUF and GISMO-mis.

A. Across all cohorts
B. Across collection method. A greater proportion of blood samples were carriers for CNVs compared to saliva samples (*OR = 1.1201, p = 0.0587*; *OR = 1.4974, p = 2.45e-13* for LOEUF and GISMO-mis, respectively)
C. Control samples across collection method. We observe a greater proportion of blood sample CNV carriers compared to saliva sample CNV carriers (*OR = 1.2001, p = 0.1243; OR = 1.2625, p = 0.0446* for LOEUF and GISMO-mis, respectively).

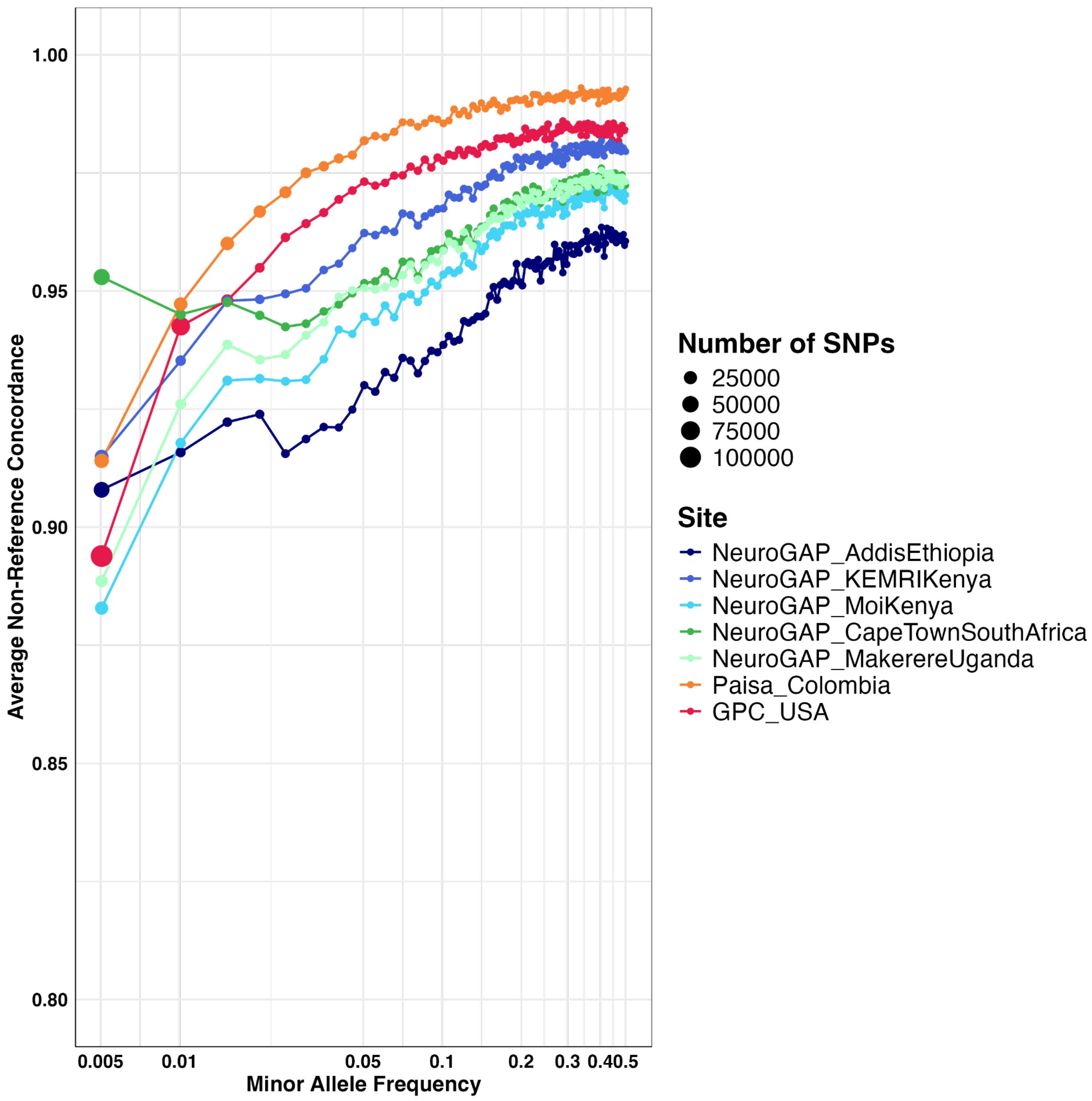


**Supplementary Figure 3.** Per-cohort non-reference concordance per minor allele frequency bin.

X-axis provides the minor allele frequency bin while the y-axis provides the non-reference concordance per cohort. Size of the points corresponds to the number of SNPs in the MAF bin. Variants are filtered to those passing an INFO score >= 0.8. MAFs per SNP are defined by the GSA array for the Paisa and GPC, given the sufficient sample sizes. MAFs are defined by the HGDP-1KG AFR subset for the NeuroGAP cohorts.

A.

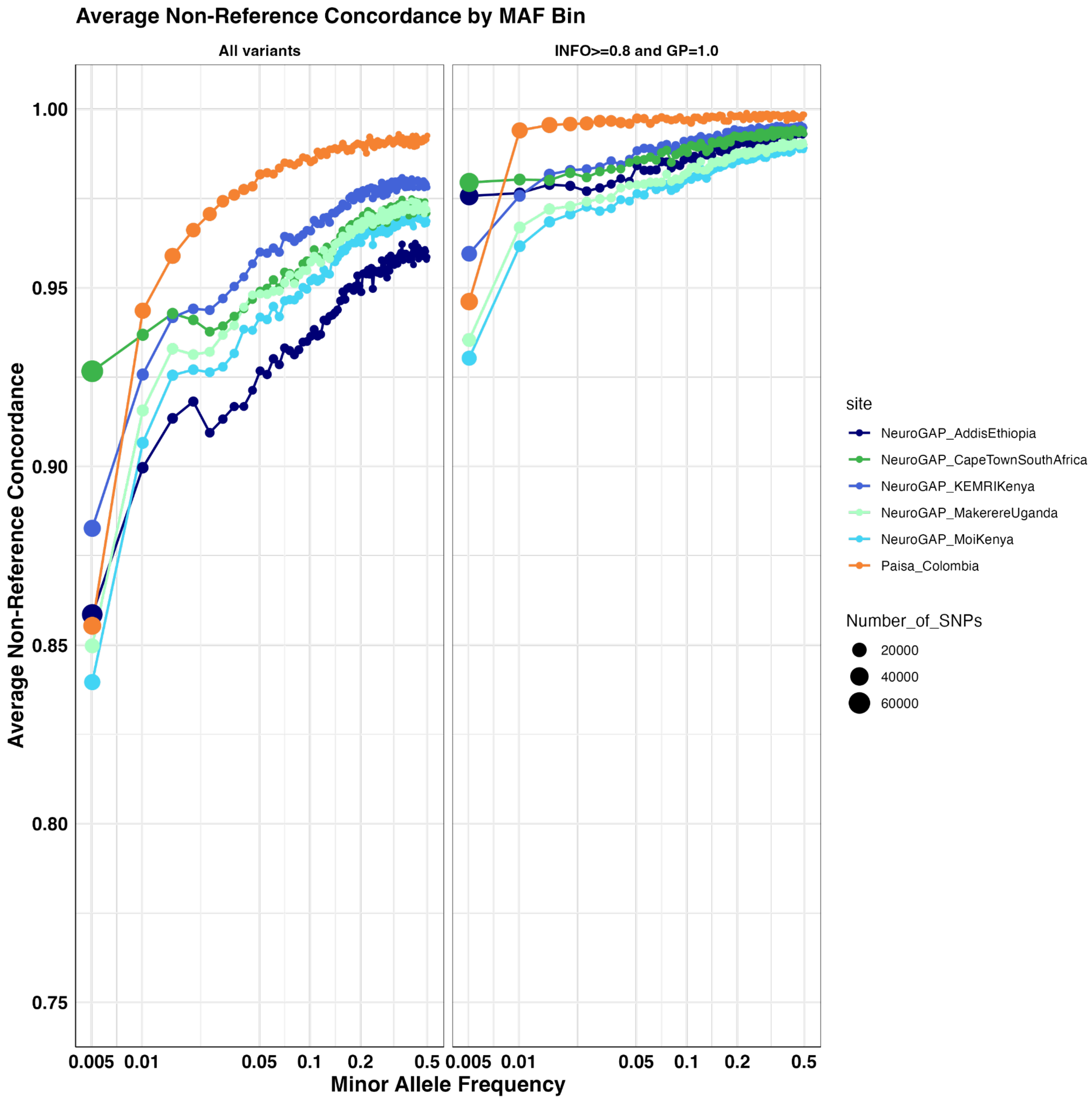

Average Non-Reference Concordance by MAF Bin
All variants
INFO>=0.8 and GP=1.0
Average Non-Reference Concordance
1.00
0.95
0.90
0.85
0.80
0.75
0.005 0.01 0.05 0.1 0.2 0.5
Minor Allele Frequency
site
NeuroGAP_AddisEthiopia
NeuroGAP_CapeTownSouthAfrica
NeuroGAP_KEMRIKenya
NeuroGAP_MakerereUganda
NeuroGAP_MoiKenya
Paisa_Colombia
Number_of_SNPs
20000
40000
60000

**B.**

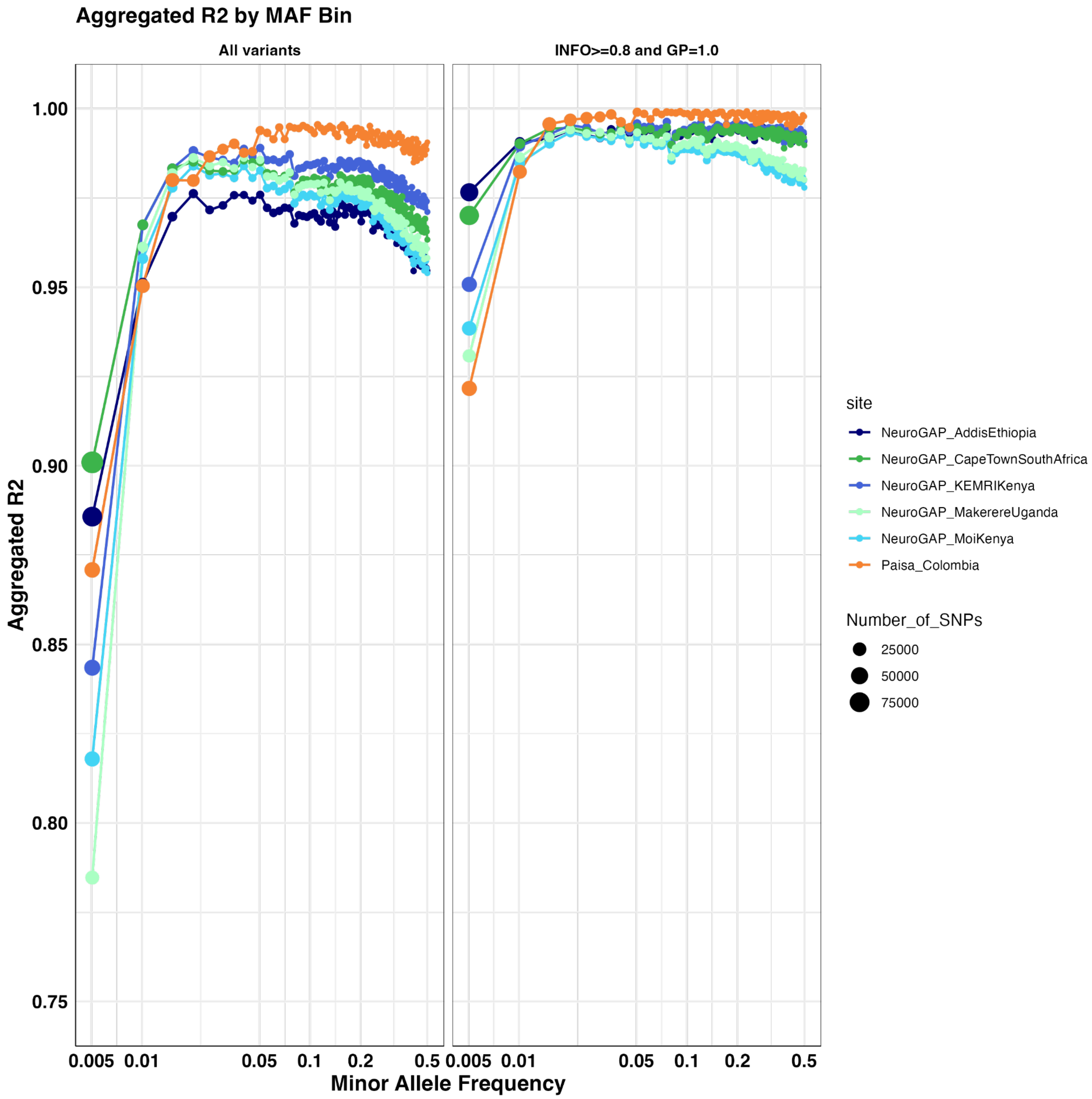

Aggregated R2 by MAF Bin
All variants
INFO>=0.8 and GP=1.0
Aggregated R2
1.00
0.95
0.90
0.85
0.80
0.75
0.005 0.01 0.05 0.1 0.2 0.5
0.005 0.01 0.05 0.1 0.2 0.5
Minor Allele Frequency
site
NeuroGAP_AddisEthiopia
NeuroGAP_CapeTownSouthAfrica
NeuroGAP_KEMRIKenya
NeuroGAP_MakerereUganda
NeuroGAP_MoiKenya
Paisa_Colombia
Number_of_SNPs
25000
50000
75000

**C.**

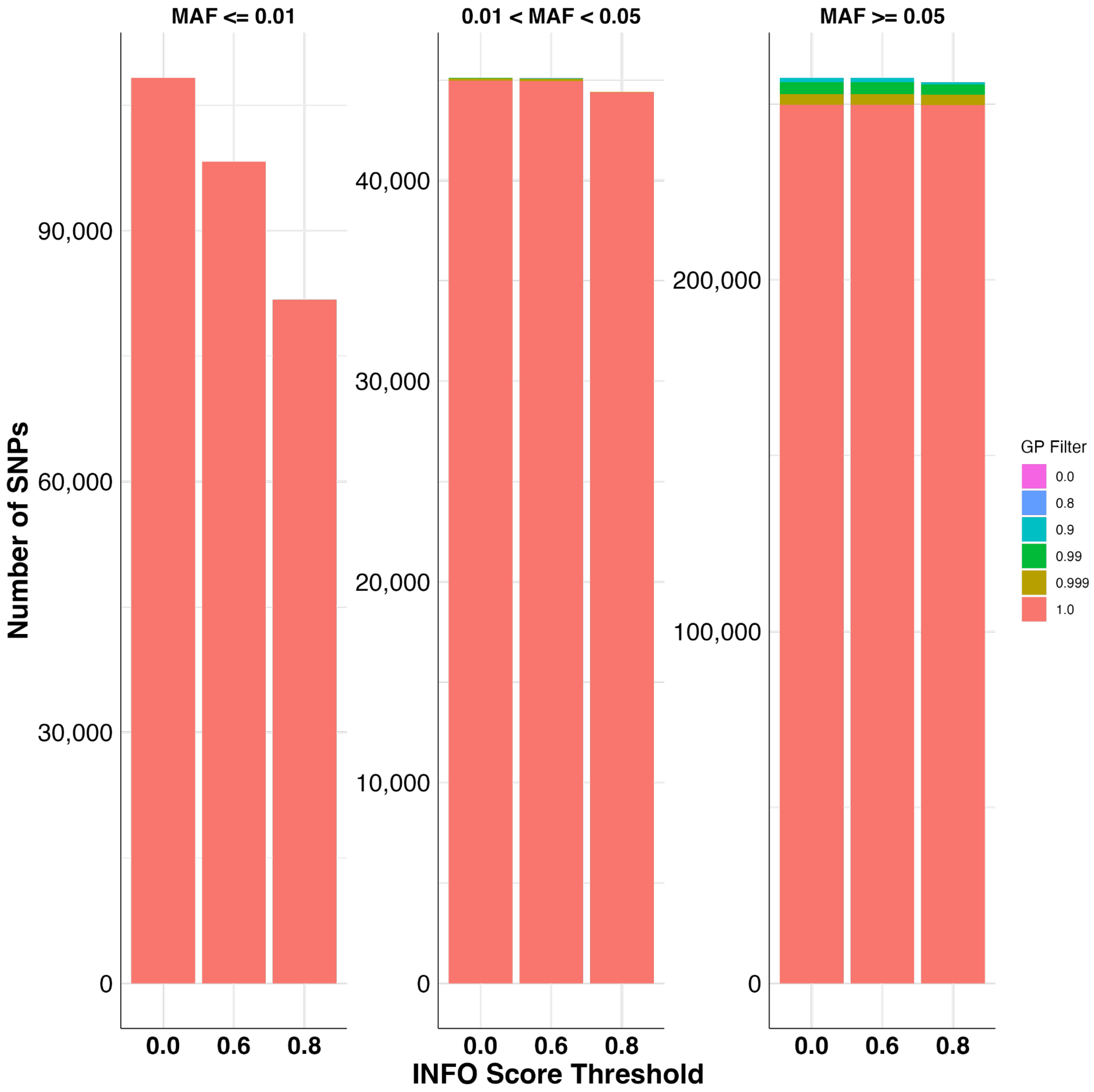


**Supplementary Figure 4.** Pre and post-QC imputation concordance and number of variants intersecting across the Illumina GSA array and imputed BGE data.

A. Non-reference concordance for the NeuroGAP and Paisa cohorts using unfiltered imputed data compared to filtering variants with INFO score at least 0.8 and requiring samples to have genotype probability (GP) = 1.0.

B. Aggregate $R^2$ for the NeuroGAP and Paisa cohorts using unfiltered imputed data compared to filtering variants with INFO score at least 0.8 and requiring samples to have genotype probability (GP) = 1.0.

C. Number of SNPs per INFO score filter and GP filter. SNPs were considered dropped by the GP filter if more than 50% of the samples did not reach the given threshold.

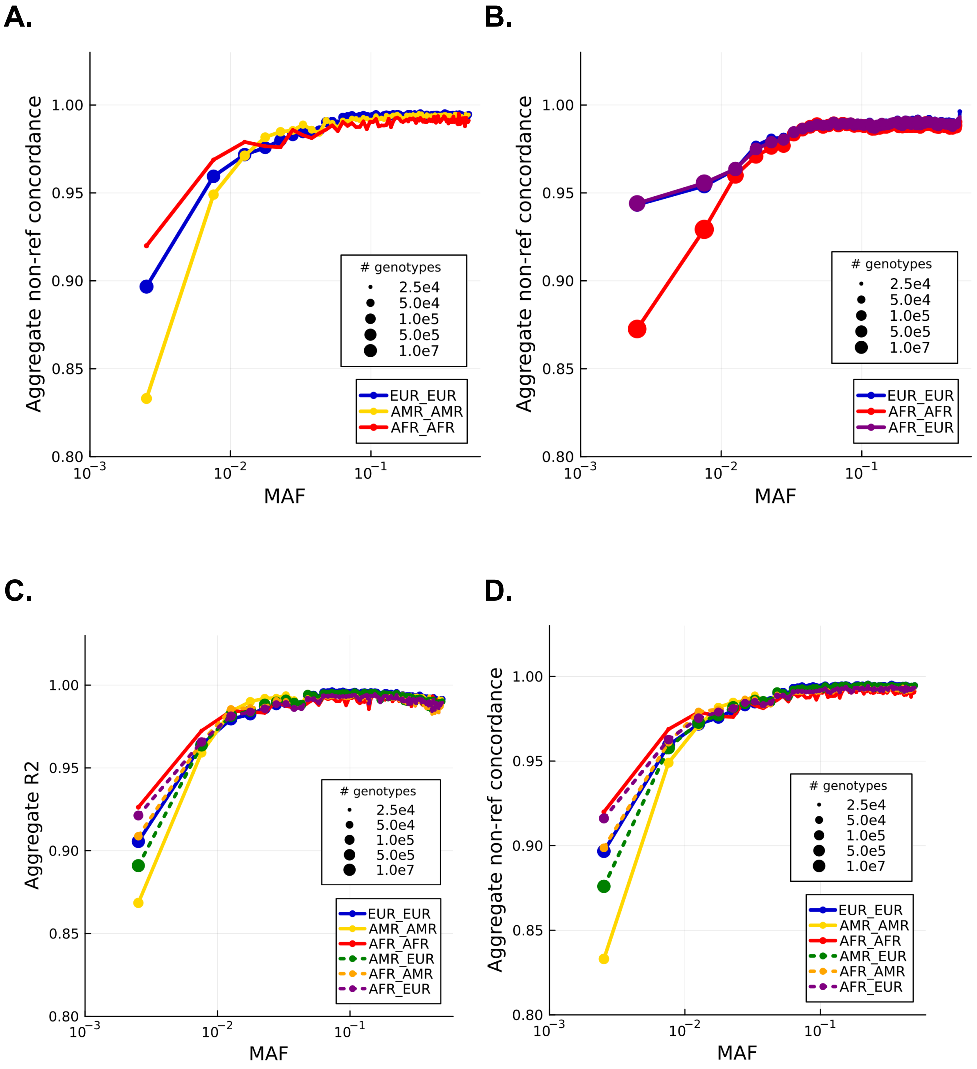


**Supplementary Figure 5.** Local ancestry imputation concordance

A. Non-reference concordance for Paisa cohort.
B. Non-reference concordance for GPC cohort.
C. Aggregated $R^2$ for Paisa cohort, including heterozygous diploid ancestry (dashed lines).
D. Non-reference concordance for Paisa cohort, including heterozygous diploid ancestry (dashed lines).
